# MLX phosphorylation stabilizes the ChREBP-MLX heterotetramer on tandem E-boxes to control carbohydrate and lipid metabolism

**DOI:** 10.1101/2024.09.04.611172

**Authors:** Carla E Cadena del Castillo, Onur Deniz, Femke van Geest, Lore Rosseels, Ingrid Stockmans, Marius Robciuc, Sebastien Carpentier, Bettina K. Wölnerhanssen, Anne Christin Meyer-Gerspach, Ralph Peterli, Ville Hietakangas, Mitsugu Shimobayashi

**Affiliations:** Clinical and Experimental Endocrinology, Department of Chronic Diseases and Metabolism, KU Leuven, Leuven, Belgium; Institute of Biotechnology, Helsinki Institute of Life Science, University of Helsinki, Helsinki, Finland; Facility for Systems Biology Based Mass Spectrometry, KU Leuven, Leuven, Belgium; St. Clara Research Ltd, St. Claraspital, Basel, Switzerland; University of Basel, Basel, Switzerland; Clarunis, University Digestive Health Care Center, St. Clara Hospital and University Hospital Basel, Switzerland

**Keywords:** Carbohydrate and lipid metabolism, fly development, sugar tolerance, ChoRE, MLX, ChREBP, CK2

## Abstract

The heterodimeric ChREBP-MLX transcription factor complex is a key mediator that couples intracellular sugar levels to carbohydrate and lipid metabolism. To promote the expression of target genes, two ChREBP-MLX heterodimers form a heterotetramer to bind a tandem element with two adjacent E-boxes, called Carbohydrate Responsive Element (ChoRE). How the ChREBP-MLX hetero-tetramerization is achieved and regulated, remains poorly understood. Here we show that MLX phosphorylation on an evolutionarily conserved motif is necessary for the heterotetramer formation on the ChoRE and the transcriptional activity of the ChREBP-MLX complex. We identified CK2 and GSK3 as MLX kinases that coordinately phosphorylate MLX. High intracellular glucose-6-phosphate accumulation inhibits MLX phosphorylation and heterotetramer formation on the ChoRE, impairing ChREBP-MLX activity. Physiologically, MLX phosphorylation is necessary in *Drosophila* to maintain sugar tolerance and lipid homeostasis. Our findings suggest that MLX phosphorylation is a key mechanism for the ChREBP-MLX heterotetramer formation to regulate carbohydrate and lipid metabolism.

## Introduction

Transcriptional regulation plays a central role for organisms to adapt their metabolism in response to nutrient availability. The basic helix-loop-helix/leucine zipper (bHLH/LZ) Mondo family transcription factors (TFs) promote the expression of carbohydrate-induced genes. In mammals, there are two Mondo paralogs, Carbohydrate Responsive Element Binding Protein (ChREBP) and MondoA^1, 2^. ChREBP controls the expression of genes encoding glycolytic and lipogenic enzymes in the liver and adipose tissue^3^. Previous studies suggested that the glucose metabolite glucose-6-phosphate (G6P) promotes ChREBP transcriptional activity^4–6^, although the underlying molecular mechanism is still unclear. To promote the expression of its target genes, ChREBP forms a complex with Max-like protein X (MLX)^7–11^. ChREBP-MLX complex specifically binds to two tandem E-box motifs separated by 5 nucleotides, defined as the Carbohydrate Response Element (ChoRE)^12^. Endogenous E-boxes in the ChoRE sequence do not perfectly match the canonical tandem E-box sequence (CACGTGxxxxxCACGTG)^13^, resulting in a weak recognition by the ChREBP-MLX complex. To overcome this weak recognition and promote the expression of the target genes, two ChREBP-MLX heterodimers need to coordinately bind to the naturally occurring imperfect E-boxes and form a heterotetramer^9^. It was demonstrated that the loop region of the HLH domain of MLX is critical for the ChREBP-MLX heterotetramer formation on the ChoRE^9^. However, the regulation and physiological role of the tetramer formation remain poorly understood. Here we uncovered an evolutionarily conserved signaling pathway that controls the ChREBP-MLX heterotetramer formation on the ChoRE, ChREBP-MLX activity, and thereby sugar responsive transcriptional regulation.

## Results

### MLX phosphorylation is necessary for ChREBP-MLX transcriptional activity

Since protein phosphorylation frequently regulates TF activity^14^, we explored whether phosphorylation of MLX regulates the ChREBP-MLX function. MLX appeared in multiple bands in immunoblots of protein lysates from mouse and human white adipose tissue (WAT), with slow-migrating bands sensitive to phosphatase treatment (**Fig. 1a** and **1b**). The faster-migrating bands in the phosphatase-treated lysates correspond to MLX isoforms, MLX-beta and MLX-gamma (**Fig. 1a** and **b**)^15^, suggesting that MLX is phosphorylated in mammalian WAT.

**Fig. 1.**
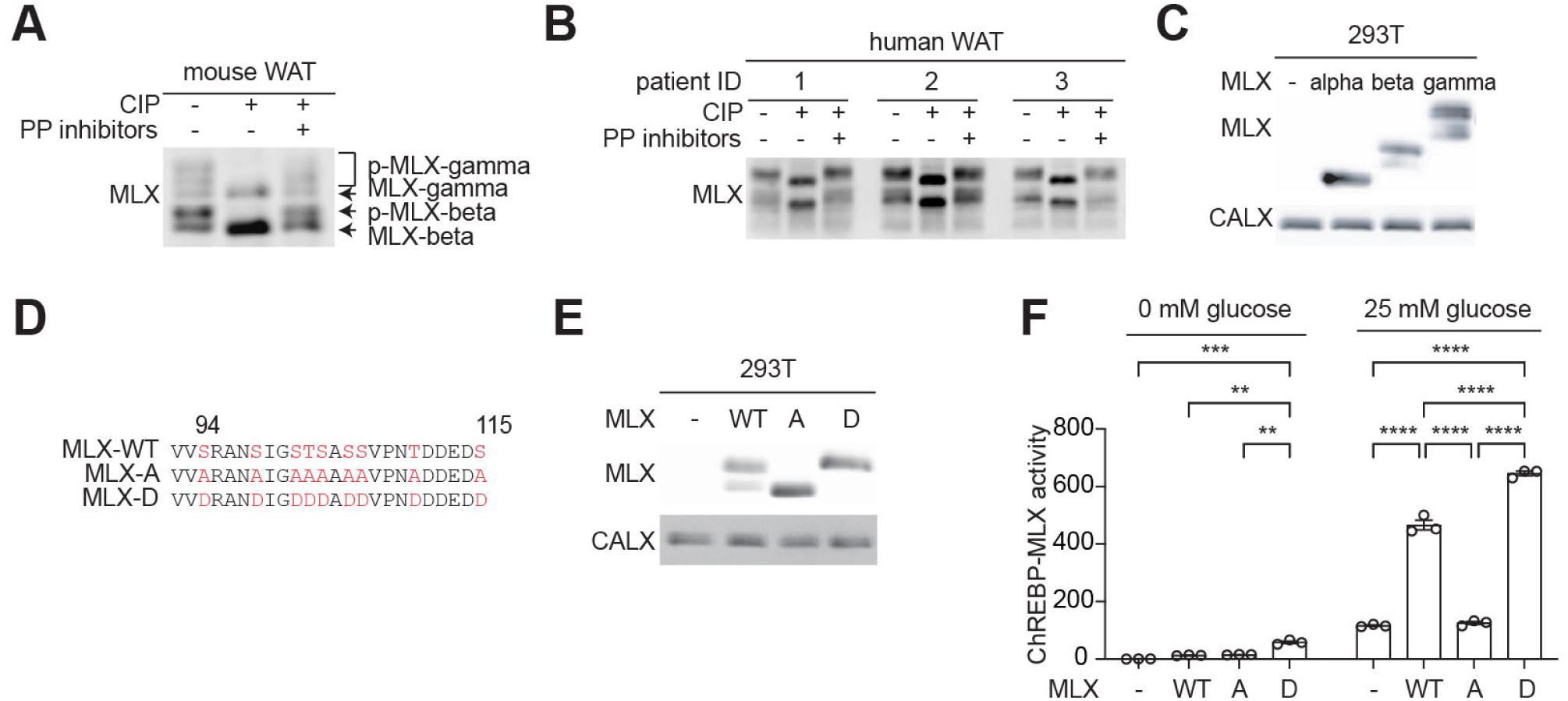
MLX phosphorylation promotes ChREBP-MLX activity. **a-b**. MLX phosphorylation in mouse (a) and human (b) white adipose tissue (WAT) lysates treated with calf intestine phosphatase (CIP) or CIP and phosphatase (PP) inhibitors. n=3. **c.** MLX phosphorylation in 293T cells expressing MLX isoforms alpha, beta, or gamma. CALX serves as a loading control. N=3. **d.** Amino acid sequence alignment of wild-type MLX (MLX-WT), phospho-deficient MLX-A or phospho-mimetic MLX-D. **e.** MLX phosphorylation in 293T cells expressing MLX-WT, -A, or -D. N=3. **f.** ChREBP-MLX luciferase reporter activity in 293T cells expressing ChREBP, HK2 and MLX-WT, -A, or -D. Cells were starved for glucose overnight and/or treated with 25 mM glucose for 3 hours. Two-way ANOVA **p<0.01, ***p<0.001, ****p<0.0001. N=3.

To investigate the role and regulation of MLX phosphorylation, we identified the phosphorylation sites on MLX. Phospho-proteomics studies have identified 22 putative serine (S), threonine (T), and tyrosine (Y) phosphorylation sites in four regions on MLX (**Fig. S1a**)^16^. To narrow down the phosphorylation sites responsible for the band shift, we analyzed MLX phosphorylation in 293T cells transiently expressing MLX-alpha, -beta, or - gamma. MLX-beta and -gamma, but not MLX-alpha, displayed slower migrating bands (**Fig. 1c**), suggesting that MLX phosphorylation detected by the band shift is specific to the regions common to MLX-beta and -gamma (P80 to D111). The mutation of S94, S98, S101, T102, S103, S105, S106, T110, and S115 to alanine (MLX-A) or aspartic acid (MLX-D) (**Fig. 1d**) prevented and mimicked MLX phosphorylation, respectively (**Fig. 1e** and **Fig. S1b**). These data show that mammalian MLX is phosphorylated on the S/T residues between S94 and S115.

To study the role of MLX phosphorylation, we examined ChREBP-MLX transcriptional activity by a reporter system^3–5, 9, 17^ (**Fig. S1c**). We optimized the reporter system by co-expressing Hexokinase 2 (HK2) to robustly monitor ChREBP-MLX activity^4, 5, 18^ (**Fig. S1d** and **1e**). We observed that MLX-A and MLX-D, compared to MLX-WT, displayed reduced and increased transcriptional activity, respectively (**Fig. 1f**). These data suggest that MLX phosphorylation is necessary for the ChREBP-MLX activity.

ChREBP is dephosphorylated and activated in response to high glucose^19^. To investigate the importance of MLX phosphorylation compared with ChREBP dephosphorylation, we examined the ChREBP-MLX transcriptional activity with a phospho-deficient ChREBP (ChREBP-A). MLX-A inhibited the transcriptional activity with ChREBP-WT and ChREBP-A (**Fig. S1f**), suggesting that MLX dephosphorylation overrides ChREBP dephosphorylation on the ChREBP-MLX regulation.

### MLX phosphorylation is required for sugar response in *Drosophila*

MLX phosphorylation sites are highly conserved across animalia (**Fig. 2a** and **Fig. S2a**). To study the physiological role of MLX phosphorylation, we used *Drosophila*, where *mlx* null mutants (*mlx*^1^) display larval sugar intolerance, reduced lipid storage, and elevated circulating sugar levels^10^. Considering the evolutionarily conservation of *Mlx*, we complemented loss of *mlx* with transgenes encoding mouse MLX-WT or MLX-A specifically in the fat body, the functional counterpart of the liver and adipose tissue (*Fb>mMlx-WT, mlx*^1^ and *Fb>mMlx-A, mlx*^1^). These transgenic flies are hereafter referred as *mMlx-WT* and *mMlx-A*. mMLX-WT, but not mMLX-A, is phosphorylated in larvae (**Fig. 2b**), confirming the functional conservation of Mlx kinases. As shown before^10^, *mlx*^1^ larvae displayed nearly normal development on a low sugar diet (LSD, 10% yeast), which was not affected by the expression of *mMlx-WT* and *mMlx-A* (**Fig. 2c** and **2d**). *mlx*^1^ larvae did not grow nor pupariate on high sugar diet (HSD, 10% yeast + 15% sucrose)^10^, but this sugar intolerance was partially rescued in *mMlx-WT* larvae (**Fig. 2d** and **2e**). Interestingly, the rescue observed in *mMlx-A* larvae was significantly weaker compared to *mMlx-WT* larvae (**Fig. 2d** and **2e**), consistent with the functional importance of Mlx phosphorylation in supporting larval development and survival on high sugar diet.

**Fig. 2.**
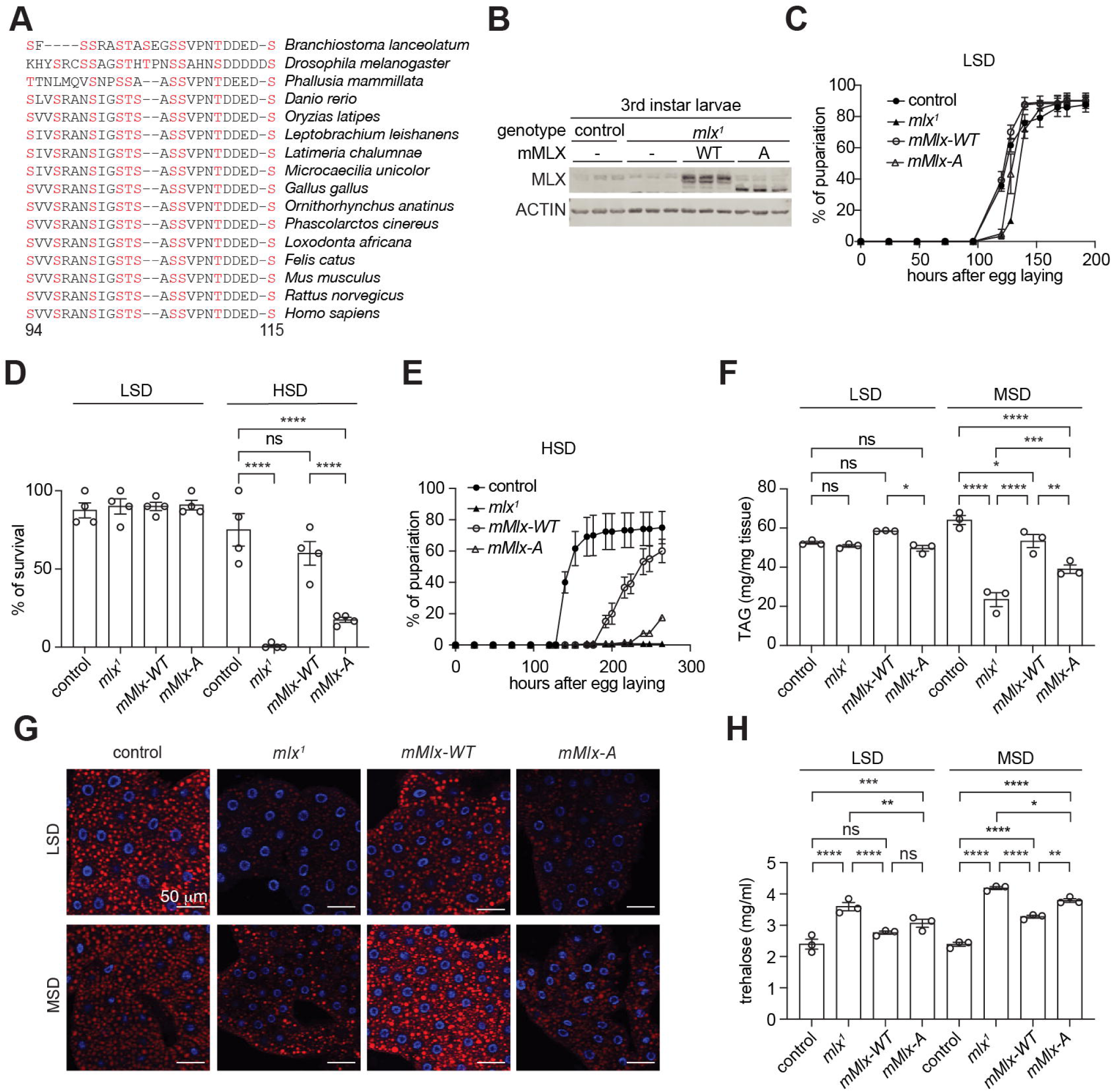
MLX phosphorylation on an evolutionarily conserved motif promotes the sugar response in *Drosophila*. **a.** Amino acid sequence alignment of MLX phosphorylation sites across different animal species. **b.** MLX phosphorylation in 3^rd^ instar control or mlx null (*mlx*^1^) larvae complemented with fat body-specific expression of mouse MLX-WT or MLX-A (*mMlx-WT* or *mMlx-A*, respectively). ACTIN serves as a loading control. n=3. **c.** Pupariation of control, *mlx*^1^, *mMlx-WT* or *mMlx-A* larvae grown in low sugar diet (LSD, 10% yeast). N=4. **d.** Survival of control, *mlx*^1^, *mMlx-WT* or *mMlx-A* larvae grown on LSD or high sugar diet (HSD, 10% yeast and 15% sucrose). Two-way ANOVA ****p<0.0001, ns=not significant. N=4. **e.** Pupariation of control, *mlx*^1^, *mMlx-WT* or *mMlx-A* larvae grown on HSD. N=4. **f.** Triglyceride (TAG) levels of 3^rd^ instar control, *mlx*^1^, *mMlx-WT* or *mMlx-A* larvae grown on LSD or medium sugar diet (MSD, 10% yeast and 5% sucrose). Two-way ANOVA *p<0.05, **p<0.01, ***p<0.001, ****p<0.0001, ns=not significant. N=3. **g.** Neutral lipid staining of the fat body in 3^rd^ instar control, *mlx*^1^, *mMlx-WT* or *mMlx-A* larvae grown in LSD or MSD. n=5. **h.** Hemolymph trehalose levels in 3^rd^ instar control, *mlx*^1^, *mMlx-WT* or *mMlx-A* larvae in LSD or MSD. Two-way ANOVA *p<0.05, **p<0.01, ***p<0.001, ****p<0.0001, ns=not significant. N=3.

*Drosophila* Mlx, together with the ChREBP ortholog Mondo, has a conserved role promoting lipogenesis in response to sugar feeding^20^. Therefore, we next examined triglyceride (TAG) levels in 3^rd^ instar larvae grown either on LSD or a medium sugar diet (MSD, 10% yeast + 5% sucrose), which unlike HSD, allows larval development of *mlx*^1^ mutants. In larvae grown on LSD, we observed slight reduction of whole-body TAG levels in *mMlx-A* larvae compared to *mMlx-WT*. (**Fig. 2f**). Compared to controls, *mlx*^1^ larvae grown on MSD displayed reduced TAG levels, which were rescued by fat body-specific expression of mMLX-WT (**Fig. 2f**). Notably, the rescue of TAG levels was significantly compromised by fat body-specific expression of mMLX-A (**Fig. 2f**). To analyze fat body lipid storage, we stained neutral lipids of 3^rd^ instar larvae with LipidTOX^TM^ (**Fig. 2g**). Both *mlx*^1^ and *mMlx-A* larvae displayed reduced lipid droplet volume in both LSD and MSD conditions, compared to control and *mMlx-WT* larvae (**Fig. 2g** and **Fig. S2b**). Fat body cells in *mlx*^1^ and *mMLX-A* were smaller than control and *mMlx-WT* (**Fig. 2g**). Reduced lipid volume was observed in *mlx*^1^ and *mMlx-A* larvae, even after normalization with cell size (**Fig. S2c**).

Impaired Mondo/ChREBP-MLX function in the fat body in flies and adipose tissue in mice causes high circulating sugar levels^10, 18, 20, 21^. Insect hemolymph contains low levels of diet-derived free glucose and much higher levels of non-reducing disaccharide trehalose, synthesized by the fat body^22, 23^. Thus, we examined trehalose and glucose levels in the hemolymph of larvae fed with LSD or MSD. Compared to control larvae, hemolymph trehalose levels were elevated in *mlx*^1^ larvae in both LSD- and MSD-fed conditions (**Fig. 2h**). *mMlx-WT* expression partially rescued the hemolymph trehalose levels, while the rescue was significantly blunted in the phospho-deficient mutant line in a MSD-fed condition. Hemolymph glucose levels were not significantly different in LSD likely due to low abundance and high variability (**Fig. S2d**). However, *mlx*^1^ and *mMlx-A* larvae had significantly higher hemolymph glucose levels, compared to control larvae in MSD (**Fig. S2d**). Overall, the data from *Drosophila* indicate that MLX phosphorylation has a physiological role in maintaining organismal sugar tolerance and controlling lipid and carbohydrate homeostasis.

### MLX phosphorylation stabilizes the heterotetrameric ChREBP-MLX complex on ChoRE

Having established the important role of MLX phosphorylation in the transcriptional activity and sugar response, we investigated whether MLX phosphorylation controls the ChREBP-MLX activity via ChoRE binding. First, we modeled two ChREBP-MLX complexes with the ChoRE sequence from the promoter region of the target gene encoding pyruvate kinase (PK) by AlphaFold 3^24^. The model showed that the MLX phosphorylation motif lies in a flexible region close to the alpha-helix which binds to the E-box (**Fig. 3a**). Interestingly, a model with phosphorylated MLX showed a drastic conformational change in ChREBP-MLX dimers, stabilizing the flexible region as alpha-helixes (**Fig. 3b**). Furthermore, the model predicted that MLX phosphorylation promotes interdimer interactions of the two ChREBP-MLX dimers, although the confidence score for this interdimer region is very low (**Fig. S3a**). These models suggest that MLX phosphorylation stabilizes the ChREBP-MLX tetramer on the ChoRE sequence.

**Fig. 3.**
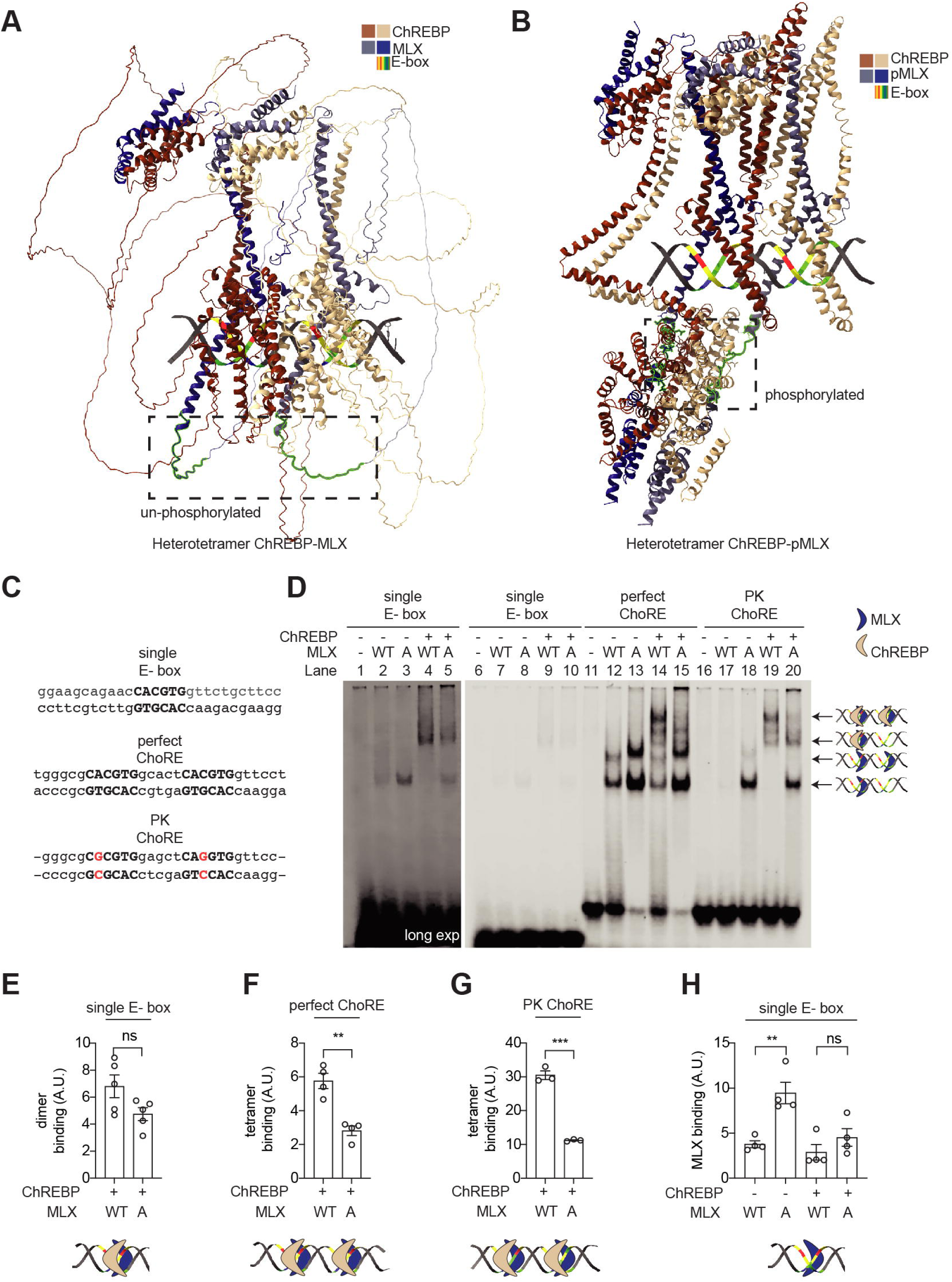
MLX phosphorylation is required for the binding of ChREBP-MLX heterotetrametric complex on the ChoRE. **a-b**. A model of ChREBP-MLX (a) and ChREBP-pMLX (b) structure on the PK ChoRE by AlphaFold 3. **c**. Sequences of single E-box, perfect-ChoRE or PK-ChoRE probes. **d**. Binding of ChREBP and/or MLX to single E-box, perfect-ChoRE or PK-ChoRE probes. **e-h**. Quantification in d. One-way ANOVA or t-test, **p<0.01***p<0.001 ns=not significant. N=3-4.

To experimentally validate the model, we examined the binding affinity of purified ChREBP and MLX to a single perfect E-box, a perfect ChoRE (two tandem perfect E-boxes), and the PK ChoRE sequence by electrophoretic mobility shift assay (EMSA) (**Fig. 3c, 3d, Fig. S3b,** and **3c**). The binding of ChREBP-MLX and ChREBP-MLX-A heterodimer to the single E-box, perfect ChoRE, and PK ChoRE was similar (**Fig. 3d**: lanes 4 vs 5, 14 vs 15, 19 vs 20 and **Fig. 3e, Fig. S3d,** and **S3e**), suggesting that MLX phosphorylation is not required for the MLX-ChREBP dimer formation and the recognition of an E-box by a ChREBP-MLX dimer. Intriguingly, the binding of ChREBP-MLX heterotetramer to the perfect ChoRE and PK ChoRE was strongly impaired in the absence of MLX phosphorylation (**Fig. 3d**: lanes 14 vs 15, 19 vs 20 and **Fig. 3f**, and **3g**). These data validate the AlphaFold 3 model and suggest that MLX phosphorylation is required for the stability of the ChREBP-MLX heterotetramer binding to the ChoRE.

Notably, MLX-A displayed higher ChREBP-independent affinity to the single E-box, perfect ChoRE, and PK ChoRE than MLX-WT (**Fig. 3d**: lanes 2 vs 3, 12 vs 13, 17 vs 18 and **Fig. 3h, Fig. S3f,** and **S3g**). It remains to be determined how the ChREBP-independent MLX E-box binding is achieved.

### CK2 and GSK3 sequentially phosphorylate MLX to promote ChREBP-MLX activity

Next, we determined the upstream signaling pathway controlling MLX phosphorylation and the heterotetramer formation. Mammalian target of rapamycin complex 1 and 2 (mTORC1/2) are protein kinase complexes that promote ChREBP activity and *de novo* lipogenesis (DNL)^25–27^, but mTOR inhibitors (rapamycin and torin2) had no impact on MLX phosphorylation (**Fig. S4a**). To identify MLX kinases, we performed an unbiased and proteomics-based proximity ligation assay^28^. By transiently expressing the functional MLX-TurboID fusion protein (**Fig. S4b** and **S4c**) in 293T cells, we searched for putative MLX kinases by mass spectrometry. Among the enriched biotinylated kinases, CSNK2A1 and CSNK2A2, catalytic subunits of CK2, were one of the top candidates (**Fig. 4a**). CK2 is known to phosphorylate the pS/T-D/E-X-D/E motif^29^, found in the MLX phosphorylation site (**Fig. 4b**). An AlphaFold 3^24^ model indeed predicts that the CK2 catalytic site binds to the putative CK2 motif on MLX (**Fig. 4c**). Consistently, the ATP-competitive CK2 inhibitor CX-4945^30^ inhibited MLX phosphorylation (**Fig. 4d**), suggesting that CK2 is an MLX kinase. CK2 has been shown to act as a priming kinase for GSK3^31, 32^. The N-terminal region of the CK2 motif on MLX phosphorylation sites contains the consensus motif of GSK3 (pS/T-X-X-X-pS/T) (**Fig. 4b**). Indeed, the GSK3 inhibitor CHIR99021 (**Fig. 4e**)^33^ or genetic depletion of GSK3 partially blocked MLX phosphorylation (**Fig. S4d** and **S4e**). These data suggest that CK2 and GSK3 are upstream kinases controlling MLX phosphorylation.

**Fig. 4.**
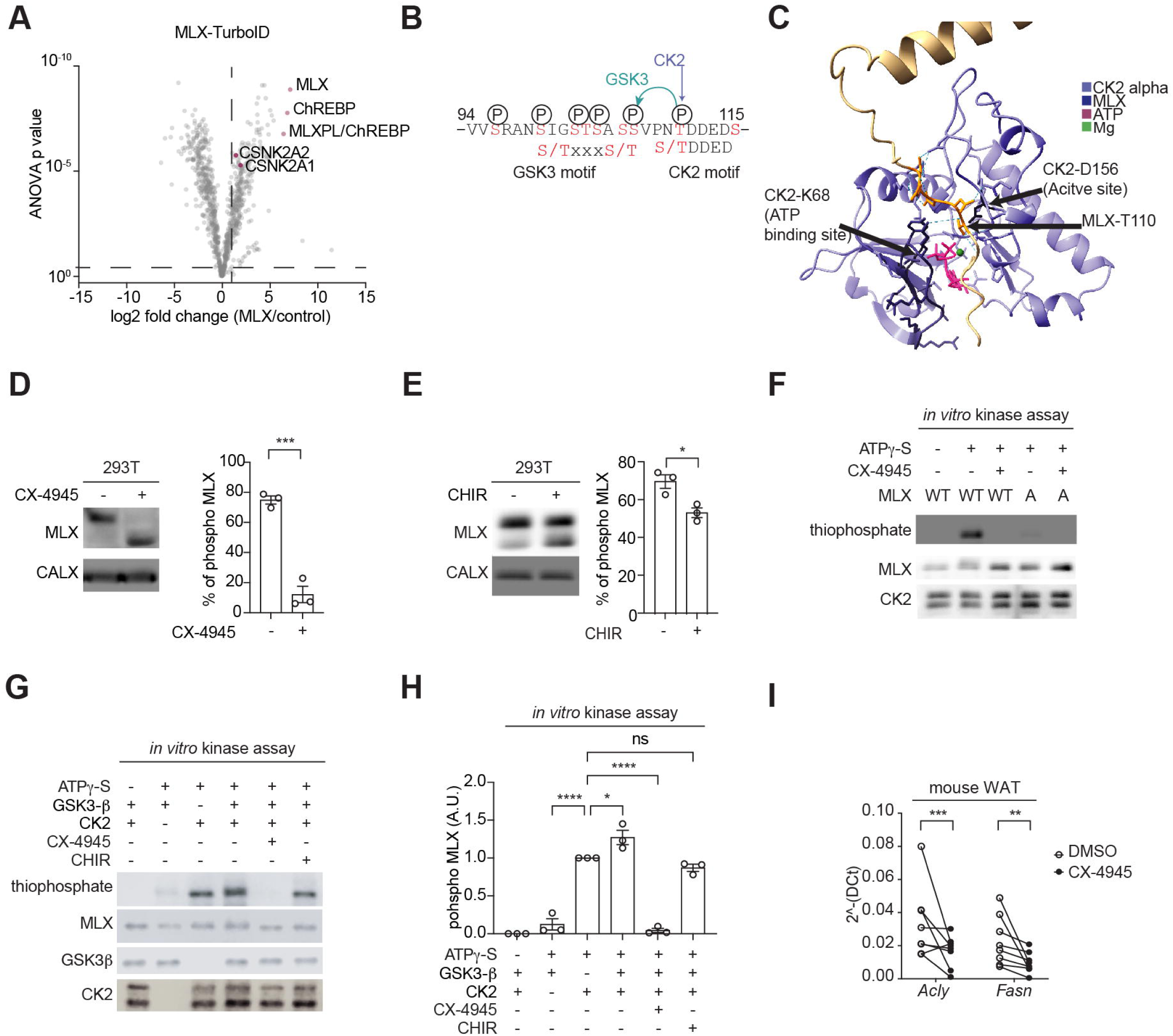
Identification of CK2 and GSK3 as MLX kinases. **a.** Interactors of TurboID-tagged MLX. 293T cells expressing ChREBP, HK2, and MLX-TurboID were labeled with 25 nM biotin for 10 min. N=4. **b.** Putative CK2 and GSK3 phosphorylation motifs on MLX. **c.** Interaction of MLX and CK2 catalytic site based on an AlphaFold3 structure prediction. **d.** MLX phosphorylation in 293T cells treated with or without the CK2 inhibitor CX-4945 (10 µM) for 60 min. CALX serves as a loading control. t-test, ***p<0.001. N=3. **e.** MLX phosphorylation in 293T cells treated with or without the GSK3 inhibitor CHIR99021 (5 µM) for 90 min. t-test, *p<0.05. N=3. **f.** *In vitro* CK2-mediated MLX phosphorylation (thiophosphate). Recombinant CK2, MLX-WT or -A, and ATPg-S were incubated with or without 25 nM CX-4945 for 30 min. N=3. **g-h**. *In vitro* CK2 and GSK3-mediated MLX phosphorylation (thiophosphate). Recombinant CK2, GSK3, MLX-WT, and ATPg-S were incubated with or without 25 nM CX-4945 or 5 µM CHIR99021 for 30 min. One-way ANOVA, *p<0.05, ****p<0.0001, ns=not significant. N=3. **i**. *Acly* and *Fasn* mRNA levels in mouse WAT explants treated with 10 µM CX-4945 for 60 min. t-test, **p<0.01, ***p<0.001. n=5.

To test whether CK2 and GSK3 directly phosphorylate MLX, we performed an *in vitro* CK2 and GSK3 kinase assay with recombinant MLX and ATP-γ-S as substrates^34, 35^. CK2 phosphorylated recombinant MLX-WT, but not MLX-A (**Fig. 4f**). Although GSK3 alone could not phosphorylate MLX (**Fig. 4g** and **4h**), GSK3 in combination with CK2 increased MLX phosphorylation. The CK2/GSK3-mediated MLX phosphorylation was completely blocked by CX-4945, but only partially blocked by CHIR99021(**Fig. 4g** and **4h**). Based on these data, we propose that CK2 phosphorylates MLX which primes GSK3-mediated MLX phosphorylation (**Fig. 4b**). Consistent with this, mutation of T110 and S115 in MLX to alanine was sufficient to block glucose-induced ChREBP-MLX activity (**Fig. S4f**). To further test the role of CK2-mediated MLX phosphorylation, we treated mouse WAT explants with the CK2 inhibitor CX-4945 and examined the expression of ChREBP-MLX target genes (**Fig. 4i**). CX-4945 decreased levels of ChREBP-MLX target genes *Acly* and *Fasn* in WAT. These data suggest that CK2 and GSK3 coordinately phosphorylate MLX, promoting ChREBP-MLX transcriptional activity.

### High G6P levels inhibit CK2-mediated MLX phosphorylation and the heterotetramer formation

Since ChREBP-MLX activity is controlled by glucose^3^, we wondered whether CK2/GSK3-mediated MLX phosphorylation correlates with glucose availability. As serum contains glucose, we examined MLX phosphorylation upon serum and glucose starvation. To our surprise, MLX phosphorylation increased in serum- and glucose-starved cells (**Fig. 5a**). Conversely, serum and glucose addition caused partial dephosphorylation of MLX (**Fig. 5b** and **5c**). Consistent with these data in cultured cells, MLX was phosphorylated in WAT of fasted mice, and MLX phosphorylation decreased upon refeeding (**Fig. 5d** and **5e**). These data suggest that MLX phosphorylation is inversely regulated by glucose availability.

**Fig. 5.**
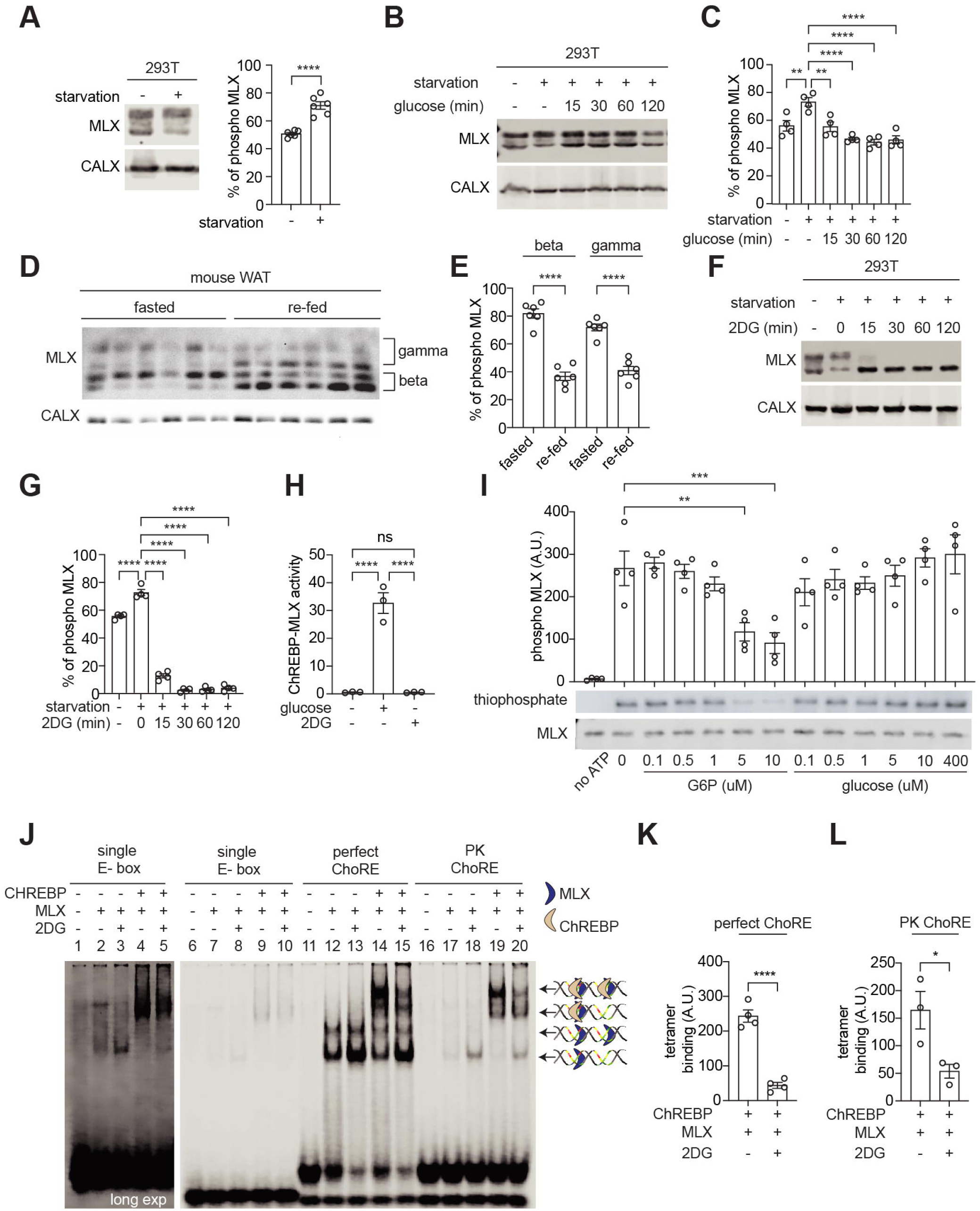
G6P accumulation inhibits CK2-mediated MLX phosphorylation and the binding of ChREBP-MLX tetramer on the ChoRE. **a.** MLX phosphorylation in 293T cells starved for glucose and serum. CALX serves as a loading control. t-test, ****p<0.0001. N=6. **b-c**. MLX phosphorylation in 293T cells starved for glucose and serum overnight and re-fed with 25 mM glucose. One-way ANOVA, **p<0.01, ****p<0.0001. N=4. **d-e**. MLX phosphorylation in WAT from overnight fasted or overnight fasted and 3-hour re-fed mice. One-way ANOVA, ****p<0.0001. n=6. **f-g**. MLX phosphorylation in 293T cells starved for glucose and re-fed with 25 mM 2DG. One-way ANOVA, ****p<0.0001. N=4. **h.** ChREBP-MLX luciferase reporter activity in 293T cells expressing ChREBP, HK2 and MLX-WT. Cells starved for glucose and re-fed with 25 mM glucose or 25 mM 2DG for 3 hours. One-way ANOVA ****p<0.0001, ns=not significant. N=3. **i.** *In vitro* CK2-mediated MLX phosphorylation (thiophosphate) in the presence of different concentrations of glucose-6-phosphate (G6P) or glucose. One-way ANOVA, **p<0.01,***p<0.001. N=4. **j.** Binding of ChREBP and/or MLX to single E-box, perfect-ChoRE or PK-ChoRE probes. ChREBP and MLX were purified from 293T cells treated with or without 25 mM 2DG for 1 hour. **k-l**. Quantification in j. t-test, *p<0.05, ****p<0.0001. N=3-4.

These effects of glucose on MLX phosphorylation seem paradoxical to our earlier observation that MLX phosphorylation is required for the stability and activity of ChREBP-MLX tetrameric complex and for sugar response (**Fig. 1f, 2e, 3d** and, **Fig. S1f**). How can we explain this paradox? We hypothesize that glucose-induced MLX dephosphorylation acts as a feedback mechanism to adjust ChREBP-MLX activity in response to glucose flux. To test this hypothesis, we treated cells with the glucose analog 2-deoxyglucose (2DG). Similar to glucose, 2DG is phosphorylated by hexokinases to produce the G6P analog 2DG-6-phosphate (2DG6P)^36^. Unlike G6P, cells do not metabolize 2DG6P, and thus 2DG6P accumulates within cells, which in turn blocks glucose flux^36^. 2DG inhibited MLX phosphorylation in a hexokinase-dependent manner (**Fig. 5f**, **5g**, **and Fig. S5a**). Unlike glucose (**Fig. S1d**), 2DG did not promote ChREBP-MLX activity (**Fig. 5h** and **Fig. S5b**). These data suggest that accumulation of G6P inhibits CK2/GSK3-mediated MLX phosphorylation, although glucose phosphorylation is required for the ChREBP-MLX activity (**Fig. S1e**)^4, 5, 18^. We further examined whether G6P directly inhibits CK2-mediated MLX phosphorylation by adding increasing concentrations of G6P in the *in vitro* CK2 kinase reaction. G6P, but not glucose, blocked CK2-mediated MLX phosphorylation at ≥ 5 μM, which is still the physiological G6P concentration range^37^ (**Fig. 5i**). These results suggest that high levels of G6P directly inhibit CK2-mediated MLX phosphorylation. In consistent with the phospho-deficient MLX-A mutant, MLX dephosphorylation by 2DG decreased the heterotetrametric binding of the ChREBP-MLX complex on the ChoRE sequence, but not the heterodimer binding on the single E-box (**Fig. 5j, 5k, 5l,** and **Fig. S5c-S5i**). These data suggest that G6P accumulation inhibits CK2-mediated MLX phosphorylation and thereby the formation of the ChREBP-MLX heterotetramer on the ChoRE.

## Discussion

In the eukaryotic genome, an E-box is a key cis-regulatory element controlling gene expression. E-boxes, present approximately 15 million times in the human genome^38^, are recognized by a homo or a heterodimer of the bHLH family of TFs^39^. To date, more than one hundred bHLH-containing family members have been identified in humans^40^. How is the specificity of a bHLH TF to a specific E-box determined? In addition to the dimmerization of particular bHLH proteins^41^ and DNA sequences adjacent to the E-box^42^, tandem E-boxes appear to be a key element to confer the target gene specificity for some bHLH TFs. Examples of such TFs include the heterotetramers of BMAL1-CLOCK^43^, TWIST1-E proteins^44^, and ChREBP/Mondo-MLX^9, 45^. As far as we are aware, the regulatory mechanisms controlling the formation of the above-mentioned heterotetrameric bHLH complexes on tandem E-boxes are unknown. In this study, we show that CK2/GSK3-mediated MLX phosphorylation on an evolutionarily conserved motif stabilizes the ChREBP-MLX heterotetrametric complex on the sugar-responsive tandem E-boxes (ChoRE) (**Fig. 6**). MLX phosphorylation is key for maintaining lipid storage and sugar tolerance in *Drosophila* (**Fig. 2**), highlighting the physiological importance of the ChREBP-MLX tetramer formation. We further demonstrate that G6P accumulation causes inhibition of CK2-mediated MLX phosphorylation, preventing the ChREBP-MLX tetramer formation on the ChoRE (**Fig. 5**). In summary, we propose that the CK2/GSK3-MLX signaling promotes the formation of ChREBP-MLX heterotetrametric complexes on the ChoRE, the ChREBP-MLX transcriptional activity, and thereby glucose response. To our knowledge, this is the first study revealing a signaling pathway that controls a bHLH tetrameric complex on tandem E-boxes in response to physiological cues.

**Fig. 6.**
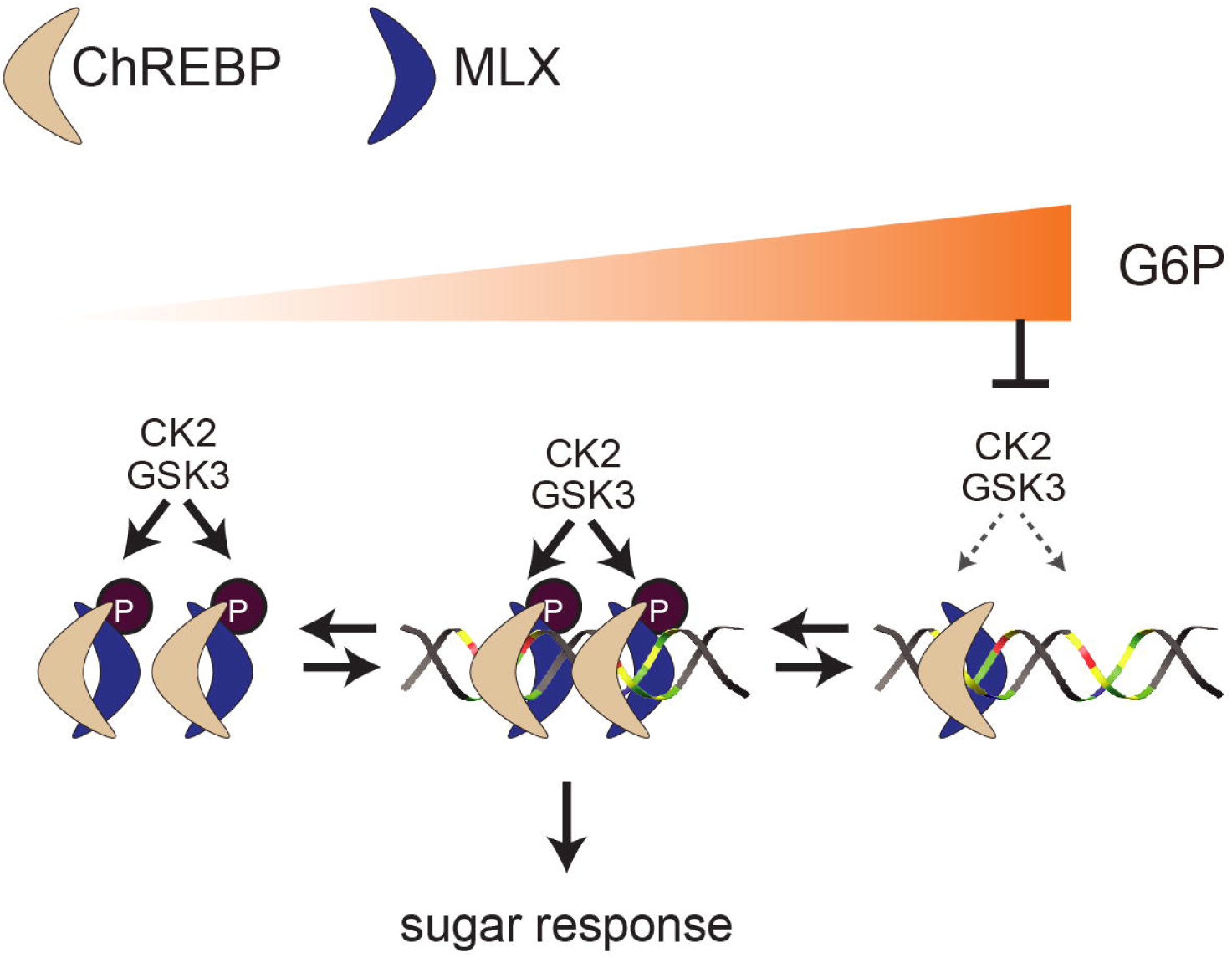
CK2/GSK3-mediated MLX phosphorylation stabilizes the heterotetrametric ChREBP-MLX complex on the ChoRE to promote sugar response. CK2 and GSK3 phosphorylate MLX in response to G6P availability. G6P accumulation inhibits MLX phosphorylation. Phosphorylated MLX stabilizes the heterotetrametric ChREBP-MLX complex on the ChoRE, promoting transcriptional activity and sugar response.

How does MLX phosphorylation promote the ChREBP-MLX heterotetramer formation on the ChoRE? A model predicted by AlphaFold 3 revealed a conformational change in both ChREBP-MLX dimers on the ChoRE (**Fig. 3b**), which favors trans-interaction between two dimers. Moreover, the model predicts that the phosphate group on S94 of human MLX from one dimer forms a hydrogen bond with human ChREBP in the adjacent dimer. Thus, MLX phosphorylation may be directly involved in the dimer-dimer interaction (**Fig. 3b**). Another possible explanation is that MLX phosphorylation promotes the interdimer interactions via the loop region of MLX since a previous model predicted that two MLX face each other and the loop region of MLX form interdimer interactions^9^. However, the model by AlphaFold 3 predicted that two ChREBP-MLX dimers bind tandem E-boxes in parallel, and we did not observe any interactions through the loop regions of MLX (**Fig. 3b**). However, when we modeled ChREBP-MLX heterotetramer with MLX loop mutant used in Ma L *et al.* ^9^, the interdimer interactions mediated by MLX phosphorylation was lost. These data may suggest that the loop region of MLX promotes interdimer interactions by bringing phosphate groups on MLX in proximity to ChREBP in the adjacent dimer. These models need to be further tested by cryo-EM structures.

Our observation that G6P inhibits CK2-mediated MLX phosphorylation seems paradoxical to the known role of G6P promoting ChREBP-MLX activity^4–6, 46^. How can G6P act as a positive and a negative regulator for the ChREBP-MLX complex? Glucose is an important carbon source for DNL through glycolysis. Glucose also provides reducing power for DNL by producing NADPH in the pentose phosphate pathway (PPP). G6P accumulation may reflect insufficient flux in glycolysis and the PPP to drive DNL. Thus, MLX dephosphorylation by G6P might serve as a feedback mechanism to couple expression of enzymes in DNL with glucose flux in glycolysis and the PPP. Alternatively, it is possible that different intercellular G6P pools activate and inhibit the ChREBP-MLX activity. Cytoplasmic G6P, directly coupled to the enzymatic activity of hexokinases, promotes the ChREBP-MLX activity, while non-cytoplasmic G6P (e.g. nuclear G6P^47^) inhibits the ChREBP-MLX tetramer formation on the ChoRE. These two hypotheses are not mutually exclusive and need to be experimentally tested.

CK2 and GSK3 are highly conserved protein kinases in eukaryotes. Since mouse MLX was phosphorylated in *Drosophila* larvae, it is likely that fly Ck2 and Sgg (fly GSK3 ortholog) phosphorylate fly Mlx. Interestingly, MLX phosphorylation sites are absent in platyhelminths (flatworms) and nematodes (roundworms). These organisms synthesize lipids in the gut and lack a specialized organ to produce lipids from carbohydrates (e.g. liver- and adipose tissue-like organs). Since MLX phosphorylation is regulated by glucose availability and is important for lipid storage, organisms with a specialized organ for lipid synthesis may have acquired CK2/GSK3-MLX signaling pathway as an evolutionary strategy to better couple carbohydrate influx to lipid metabolism. Further investigations are necessary to identify the evolutionary origin and to reveal the significance of CK2-MLX pathway across different organisms.

Our findings may shed light on pathogenesis of metabolic diseases. DNL in the liver and adipose tissue plays a crucial role in maintaining whole-body glucose homeostasis, diabetes, metabolic dysfunction-associated steatotic liver disease, and liver cancer^48–50^. Furthermore, MLX also plays a pivotal role in MYC-induced tumorigenesis^51, 52^, underscoring significance of MLX function in tumor development. Our findings of GSK3 and CK2 as MLX kinases may suggest that targeting these kinases might be effective therapeutics for the above-mentioned metabolic diseases.

## Acknowledgements

We acknowledge support from European Foundation of the Study of Diabetes/Novo Nordisk Foundation (NNF20SA0066173 to MS), KU Leuven internal fund (C14/22/116 and KAC14/22/116 to MS and PDMT1/22/015 to CECC), Fonds Wetenschappelijk Onderzoek (FWO, G023824N to MS and VH and 1297724N to CECC), Research Council of Finland (312439 to VH), Sigrid Juselius Foundation (to VH), Novo Nordisk Foundation (NNF22OC0078419 to VH). We thank for reagents and resources from Prof. Geert Carmeliet (KU Leveun), Prof. Annemieke Verstuyf (KU Leuven), Prof. Michael Schupp (Charité University), Prof. Alice Y Ting (MIT), and Prof. Feng Zhang (Broad Institute) and for technical assistance from Heini Lassila and Nicole Lamichane (University of Helsinki). We thank Kusay Arat for technical assistance for proteomics. Molecular graphics and analyses performed with UCSF ChimeraX, developed by the Resource for Biocomputing, Visualization, and Informatics at the University of California, San Francisco, with support from National Institutes of Health R01-GM129325 and the Office of Cyber Infrastructure and Computational Biology, National Institute of Allergy and Infectious Diseases. Experiments with *Drosophila* were facilitated by the University of Helsinki Drosophila core facility (Hi-Fly) and the Light microscopy unit (LMU) supported by Biocenter Finland and Helsinki Institute of Life Science.

## Author contributions

CECC, VH, and MS conceived and designed experiments, except for the *Drosophila* experiments, which were designed by VH, OD and MR in cooperation with CECC and MS. CECC, OD, FG, LR, IS, MR, and MS conducted the experiments. CECC, OD, VH and MS analyzed the data. SC supervised proteomics analysis. BKW, ACMG, and RP provided human WAT samples. VH and MS supervised the project. CECC and MS wrote the manuscript with input from all authors.

## Declaration of interests

The authors declare no competing interests.

## Methods

### Human samples

Omental white adipose tissue (oWAT) biopsies were obtained from lean subjects with normal fasting glucose level and body mass index (BMI) < 27 kg/m^2^ ^18^. All subjects gave informed consent before the surgical procedure. Patients were fasted overnight and underwent general anesthesia. All oWAT specimens were obtained between 8:30 and 12:00 am, snap-frozen in liquid nitrogen, and stored at -80 °C for subsequent use. The study protocol was approved by the Ethikkomission Nordwest-und Zentralschweiz (EKNZ, BASEC 2016-01040).

### Animals

C57BL/6 mice were purchased from an internal stock from the KU Leuven animal facility and epididymal WATs were collected. For analysis of MLX phosphorylation (Fig. 1a), WAT samples were snap-frozen and kept in -80 °C for subsequent use. For the explant experiment (Fig. 4i), WATs were washed with PBS and cultured in Dulbecco’s Modified Eagle Medium (DMEM) high glucose (Thermo Fischer Scientific, 41965062) supplemented with 1 mM sodium pyruvate (Thermo Fischer Scientific, 11360039), 1 x penicillin and streptomycin (Sigma-Aldrich, P4333), and 10% fetal bovine serum (FBS, Thermo Fischer Scientific, 10270106 lot. 2319372) and treated with 10 µM CX-4945 (Biorbyt, ORB180973) for 60 min. The tissues were washed with ice-cold PBS, snap-frozen, and kept in -80 °C for subsequent use. The study protocol was approved by KU Leuven animal ethical committee (#206/2020).

### Cell culture

293T cells were obtained from Prof. Geert Carmeliet (KU Leuven) and 3T3-L1 cells from ATCC. Cells were cultured in M1 medium composed of DMEM high glucose (Thermo Fischer Scientific, 41965062) supplemented with 1 mM sodium pyruvate (Thermo Fischer Scientific, 11360039), 1 x penicillin and streptomycin (Sigma-Aldrich, P4333), and 10% fetal bovine serum (FBS, Thermo Fischer Scientific, 10270106 lot. 2319372). Cells were cultured at 37 °C incubator with 5% CO_2_. For starvation, cells were washed twice with starvation media (DMEM no glucose, Thermo Fischer Scientific, 11966025) supplemented with 1 mM sodium pyruvate (Thermo Fischer Scientific, 11360039), 1 x penicillin and streptomycin (Sigma-Aldrich, P4333) and kept in starvation media. For GSK3 knockout/knockdown cells were generated by transfecting 293T cells with the corresponding lentiCRISPR v2 plasmid with jetPRIME (Polyplus-transfection, Cat# 101000046) and selected with puromycin (InvivoGen ant-pr-1) (1 μg/ml) for 2 days followed by transfection of GSK3-beta siRNA (Integrated DNA Technologies, Inc., Design ID: hs.Ri.GSK3B.13.1, hs.Ri.GSK3B.13.2, hs.Ri.GSK3B.13.3). For overexpression experiments in 293T cells, unless otherwise stated, cells were transfected with plasmids harboring genes encoding mouse ChREBP, rat HK2 and mouse MLX with jetPRIME (Polyplus-transfection, Cat# 101000046). 293T cells were treated with 100 nM rapamycin (LC Laboratories, R-5000 in DMSO), 5 μM CHIR99021 (Sigma-Aldrich, SML1046 in DMSO), 250 nM torin2 (LC Laboratories, T-8448 in DMSO) for 90 minutes or with 10 μM CX-4945 (Biorbyt, ORB180973 in DMSO) for 60 minutes. 3T3-L1 cells were cultured and differentiated as previously described ^18^. For differentiation, cells were maintained in M1 medium for 2 days after reaching confluence followed by two day treatment with M2 medium composed of M1 medium supplemented with 1.5 µg/mL insulin (Sigma-Aldrich, Cat# I9278), 0.5 mM IBMX (AdipoGen LIFE SCIENCES, Cat# AG-CR1-3512-G001), 1 μM dexamethasone (Sigma-Aldrich, D4902), and 2 μM rosiglitazone (AdipoGen LIFE SCIENCES, AG-CR1-3570), followed by a 4 day incubation with M3 medium (M1 with 1.5 μg/mL insulin). Differentiated cells were maintained in M3 with medium change every 2 days until used.

### Plasmids

For protein expression, cDNA was cloned into pcDNA3 for mammalian expression, pGEX for bacterial expression, or pUAST for fly experiments by using the primers listed in **Supplementary data table 1**. Mutations were introduced by reverse PCR (**Supplementary data table 1**), followed by DpnI (New England Biolabs, R0176L) digestion and bacteria transformation. For GSK3 KO sgRNAs targeting GSK3a (**Supplementary data table 1**) were cloned into lentiCRISPRv2 (a gift from Prof. Feng Zhang) as previously described ^53^. Plasmid sequences were validated.

### Immunoblots

Tissues or cells were lysed in a lysis buffer (100 mM Tris (Sigma-Aldrich, T1503) pH7.5, 2 mM EDTA (Fisher Scientific,10522965), 2 mM EGTA (Sigma-Aldrich E4378), 150 mM NaCl (Fisher Chemical S/3160/60), 1% Triton X-100 (Fisher Chemical T/3751/08), cOmplete (TM), Mini, EDTA-free Protease I (Roche, 469315900) and PhosSTOP (Roche, 4906837001). For tissue samples protein concentration was determined by Bradford assay (Bio-rad, 5000006). Equal amounts of proteins were separated by SDS-PAGE and transferred onto nitrocellulose membranes (Amersham™ Protran® Premium Western blotting membranes, GE10600003). Membranes were blocked with Interceptor reagent (Li-COR 927-70001) and antibodies were diluted with 3% BSA (Fisher Scientific, BP9703-100). The antibodies are listed in **Supplementary data table 2**. Images were captured with Li-COR Odyssey XF. Quantification was done by ImageJ software (NIH).

### CIP treatment

Protein lysates from mouse WAT or 293T cells transiently expressing ChREBP, MLX and HK2 were incubated with 5 µL of CIP (Quick CIP New England Biolabs, M0525S) and/or PhosSTOP (Roche, 4906837001) in 40 µL of 1x Cutsmart buffer (New England Biolabs, B6004S). The reaction was incubated at 37°C for 30 min with 800 rpm shaking. The reaction was stopped by adding 12.5 µL 5x SDS sample buffer, and the samples were incubated at 65 °C for 15 min before immunoblot.

### Luciferase assay

293T cells were transfected with pGL3-ChoRE-luc plasmid, pNL1.1[Nluc/TK] plasmid, and plasmids described in **Supplementary data table 3** hours post transfection, cells were starved for glucose for 16 hours and re-fed with 25 mM glucose or 2DG for 3 hours. Luciferase activity was measured with Nano-Glo® Dual-Luciferase® Reporter Assay System (Promega N1521) according to the manufacturer’s instructions.

### Fly stocks and husbandry

Flies were maintained at 25 °C in a 12LJhour light/12LJhour dark cycle, on standard *Drosophila* medium (agar 0.6% (w/v), semolina 3.2% (w/v), malt 6.5% (w/v), dry baker’s yeast 10% (w/v), propionic acid 0.7% (v/v), and Nipagin (methylparaben) 2.4% (v/v). Sucrose 5% (w/v) or 15% (w/v) was added for medium or high sugar diet, respectively. The following lines were generated in this study *UAS-mMlx-WT*, *UAS-mMlx-A*. Wild type mouse Mlx and the mutant version were cloned into the pUASTLJattB vector and directed to the attP2 landing site on chromosome 3. Transgenic flies were created by WellGenetics Inc. and recombined with the mlx-null mutant flies, *mlx*^1^ ^10^. As control, we recovered lines from which the P-element had been excised precisely, leaving *mlx* intact ^10^. The following genotypes of larvae were used in the analyses: FB-Gal4; Control, FB-Gal4; *mlx*^1^, FB-Gal4; *UAS-mMlx-WT*, *mlx*^1^, FB-Gal4; *UAS-mMlx-A*, *mlx*^1^

### Fly pupariation and growth analysis

Flies were allowed to lay eggs for 24 hours on apple juice plates (33.33% apple juice (v/v), 1.75% agar (w/v), 2.5% sugar (w/v), 2.0% Nipagin (v/v)), supplemented with dry yeast, kept at 25 °C. After 24 hours, 30 x 1st instar larvae were collected to vials containing yeast or sugar diets and kept at 25 °C. Pupariation was scored every 24 hours, and survival at the end of the observation period.

### LipidTOX staining

Fat bodies from early 3^rd^ instar larvae were fixed in 4% formaldehyde for 30 min and washed three times with PBS. Fat bodies were stained with LipidTOX (Thermo Fischer Scientific, H34477) 1:400 in PBS for 30 min at room temperature, washed three times with PBS, and mounted using VECTASHIELD Mounting Medium with DAPI (VectorLabs, H-1200). Samples were imaged using a Leica SP8 up-right microscope, and images were processed using Imaris software (Oxford Instruments).

### LipidTOX image quantification

For lipid droplet analysis, the cell borders were defined in multiple z layers using Imaris custom surface tool. Lipid staining was masked within the rendered cell surface to have lipids within cell borders. Surface detection tool was used on the masked lipid staining to get all lipid volume within the cell. The lipid volume was normalized with cell volume.

### Fly TAG analysis

For each replica 10 x early 3rd instar larvae per group were collected and snap frozen in liquid nitrogen. Larvae were homogenized in 300 μL cold 1xPBS (+0.05% Tween 20), and the lysates were inactivated at 70 °C for 10 min. Glycerol and TAG levels were measured by using the coupled colorimetric assay kits (Free Glycerol kit - Cayman Chemical; 10010755, Triglyceride kit-Cayman Chemical; 10010303). The values were normalized to body weight.

### Fly sugar analysis

Hemolymph from 3rd instar larvae was extracted as described previously ^54^ and the circulating glucose and trehalose were measured with the Glucose HK assay reagent (Sigma; GAHKLJ20) and trehalase from porcine kidney (Sigma; T8778-1U) as described previously^55^.

### BioID

293T cells were transfected with plasmids harboring genes encoding ChREBP, HK2, TurboID-tagged MLXs. 24 hours after transfection, cells were treated with 25 nM biotin for 10 min. Cells were lysed in lysis buffer (100 mM Tris (Sigma-Aldrich, T1503) pH7.5, 2 mM EDTA (Fisher Scientific,10522965), 2 mM EGTA (Sigma-Aldrich E4378), 150 mM NaCl (Fisher Chemical S/3160/60), 1% Triton X-100 (Fisher Chemical T/3751/08), cOmplete (TM), Mini, EDTA-free Protease I (Roche, 469315900) and PhosSTOP (Roche, 4906837001). Biotinylated proteins were pulled down by Streptavidin-coupled magnetic beads (Thermo Fischer Scientific) according to^28^. Proteins were precipitated by trichloroacetic acid, alkylated, digested with modified trypsin (enzyme/protein ratio 1:50) overnight. Peptides were desalted using C18 reverse-phase spin columns (Macrospin, Harvard Apparatus) according to the manufacturer’s instructions, dried under vacuum, and stored at −20 °C until further use.

### Proteomics

The digested samples were injected (5 μL) and separated on an Ultimate 3000 UPLC system (Dionex, Thermo Fischer Scientific) equipped with an Acclaim PepMap100 pre-column (C18 particle size 3 μm pore size–100 Å, diameter 0.075 mm, length 20 mm, Thermo Fischer Scientific) and a C18 PepMap RSLC (particle size 2 μm, pore size–100 Å, diameter 50 μm, length-150 mm, Thermo Fischer Scientific) using a linear gradient (0.300 μL/min). The composition of buffer A is pure water containing 0.1% Formic Acid. The composition of buffer B is pure water containing 0.08% Formic Acid and 80% Acetonitrile. The fraction of buffer B increased from 0–4% in 3 min, from 4–10% in 12 min, from 10–35% in 20 min, from 35–65% in 5 min, from 65–95% in 1 min, stayed at 95% for 10 min. The fraction of buffer B decreased from 95–5% in 1 min and stayed at 5% for 10 min. The Q Exactive Orbitrap mass spectrometer (Thermo Fischer Scientific) was operated in positive ion mode with a nanospray voltage of 2.1 kV and a source temperature of 250 °C. Pierce LTQ Velos ESI positive ion calibration mix (Thermo Fischer Scientific, 88323) was used as an external calibrant. The instrument was operated in data-dependent acquisition mode with a survey MS scan at a resolution of 70,000 (fwhm at m/z 200) for the mass range of m/z 400–1600 for precursor ions, followed by MS/MS scans of the top ten most intense peaks with +2, +3, +4, and + 5 charged ions above a threshold ion count of 1e+6 at 17,500 resolution using normalized collision energy of 25 eV with an isolation window of 3.0 m/z, Apex trigger of 5-15 sec and dynamic exclusion of 10 sec. All data were acquired with Xcalibur 3.1.66.10 software (Thermo Fischer Scientific). Peptide and protein quantities were computed with the software Progenesis® (Nonlinear Dynamics). For protein identification, the spectra were exported and submitted to the protein identification software Mascot (Version 2.2.06; Matrix science, London, England) and checked against all entries present an in house uniprot based data base where ChREBP, MLX turbo and rnHK2 were added as additional entries. Spectra were searched with a mass tolerance of 10 ppm on the precursor mass and 0.02 Da on the fragments. Tolerated variable modifications were oxidation of M and deamidation of NQ. Fixed modification: carbamidomethyl C. Tolerated miscleavages was set to 2. Mascot results were imported back in Progenesis® (Nonlinear Dynamics).

### Purification of GST-MLX fusion protein

BL21 cells (New England Biolabs, C2527H) harboring pGEX-MLX-WT or -A plasmids were cultured in LB media to OD = 0.6 at 30°C, and the expression of recombinant MLX was induced by 0.25 mM of isopropyl-β-D-thiogalactopyranoside for 5 hours. Bacteria was lysed with a lysis buffer (20 mM HEPES (VWR, 30487.297) pH7.5, 2 mM EDTA (Fisher Scientific, 10522965), 100 mM NaCl (Fisher Scientific, S/3160/60), 2 mM beta-Mercaptoethanol, cOmplete(TM), Mini, EDTA-free Protease I (Roche, 4693159001), 1% Triton X-100 (Fisher Scientific, T/3751/08)) with sonication for 4 X 30 sec on ice. GST fusion proteins were purified using glutathione agarose beads (Thermo Fischer Scientific, 16100) and eluted with glutathione (Sigma-Aldrich, G4251-10G).

### *In vitro* kinase assay

*In vitro* kinase reaction was performed in 40 μl solution consisting of 1X NEBuffer™ for Protein Kinases (PK) (New England Biolabs, B6022SVIAL), 2 mM ATP-gamma-S Kinase substrate (Abcam, ab138911), 1 unit (100nM) Casein Kinase II (CK2) (New England Biolabs, P6010SVIAL) and/or recombinant human GSK3 beta (Abcam, ab63193). After incubation at 30 °C for 30 min, 30 μl of the reaction were alkylated with 1.5 µl of 50 mM p-Nitrobenzyl mesylate and Alkylation reagent (Abcam, ab138910) for 1 hour at room temperature. The reaction was stopped by adding 6 μl of 5× SDS sample buffer, and the sample was incubated at 65 °C for 10 min. The proteins were separated by SDS-PAGE and transferred to nitrocellulose membrane for immunodetection. Membranes were blocked with Interceptor reagent (LiCOR). Phosphorylated MLX was detected with anti-Thiophosphate ester antibody (Abcam, ab92570) and secondary IRDye® 800CW Goat anti-Rabbit IgG (LI-COR Biotech 926-32211).

### RNA isolation and RT–PCR

Total RNA from mouse WAT and cultured cells was isolated with TRIzol reagent (Sigma) and NucleoSpin® RNA (M&N 740955_250) followed by cDNA synthesis using iScript cDNA synthesis kit (BioLJRad 1708891). Semiquantitative realLJtime PCR analysis was performed using fast SYBR green (Applied Biosystems 10631376) on a StepOnePlus RealLJTime PCR System (Applied Biosystems). Relative expression levels were determined by normalizing to S18 expression using the ΔΔCT method. The sequence for the primers used in this study can be found in the **Supplementary data table 1**.

### Electrophoretic Mobility Shift Assay (EMSA)

1-2 mg protein lysates of 293T cells overexpressing FLAG-tagged MLX alone or FLAG-tagged ChREBP and FLAG-tagged MLX were immunoprecipitated for 2.5 hours at 4 °C with 25 μL Anti-FLAG® M2 Affinity Gel (Sigma-Aldrich, A2220). Samples were eluted with 0.2 μg/mL 3X FLAG® Peptide (Sigma-Aldrich, F4799). Eluted proteins were used for the Odyssey® EMSA assay (829-07910). Briefly, a 20 µl reaction was set using 5 μl of purified protein lysate, 1 μl of 50 nM IRD800 5 ”end-labeled aligned DNA probe, 2 μl of 25 mM DTT, DTT/ 2.5%Tween 20, 1 µg/µL Poly (dI•dC), and 1 μl of 50% glycerol in 1 x Binding Buffer (10 mM Tris (Sigma-Aldrich, T1503) pH 7.5, 50 mM KCl, 1 mM DTT). For validation of the bound proteins, 1 μl of anti-MLX (Cat# 85570, CST) or anti-ChREBP antibodies (Cat# 58069, CST) was added. Samples were separated in a pre-runed 4.5% native polyacrylamide gel in 0.5 x TBE + 2.5% glycerol buffer at 50 V for 135 minutes. The images were captured with Li-COR Odyssey XF. Quantification was done using ImageJ software (NIH).

### AlphaFold prediction

AlphaFold3 prediction was performed with Google DeepMind using human CSNK2A1, MLX gamma sequence and ChREBP with the PK ChoRE sequence (gggcgCGCGTGgagctCAGGTGgttcc). Models were analyzed by UCSF Chimera X^56^

### Alignment

For conservation analysis MLX protein sequences and alignment were obtained from the Uniprot database (https://www.uniprot.org/ ). Phylogenic analysis was perfomed with (https://www.ebi.ac.uk/jdispatcher/phylogeny/simple_phylogeny)^57^

## Data analysis

All data are shown as the mean ± SEM. Sample numbers are indicated in each figure legend. *n* represents the independent biological replicates, and N represents the number of independent experiments. To determine the statistical significance, t-test for two comparisons, one-way ANOVA for multiple comparisons, and two-way ANOVA for group analysis were performed by GraphPad Prism 9 (GraphPad Software). *p* values are indicated on each figure.

## Materials availability

Further information and requests for resources and reagents should be directed to and will be fulfilled by the corresponding author, Mitsugu Shimobayashi.

## Data and code availability

The mass spectrometry proteomics data have been deposited to the ProteomeXchange Consortium via the PRIDE^58^ partner repository with the dataset identifier PXD054667 and 10.6019/PXD054667. This study did not generate any unpublished code, software, or algorithm. Any additional information required to reanalyze the data reported in this paper is available from the lead contact upon request.

**Supplementary data table 1.**
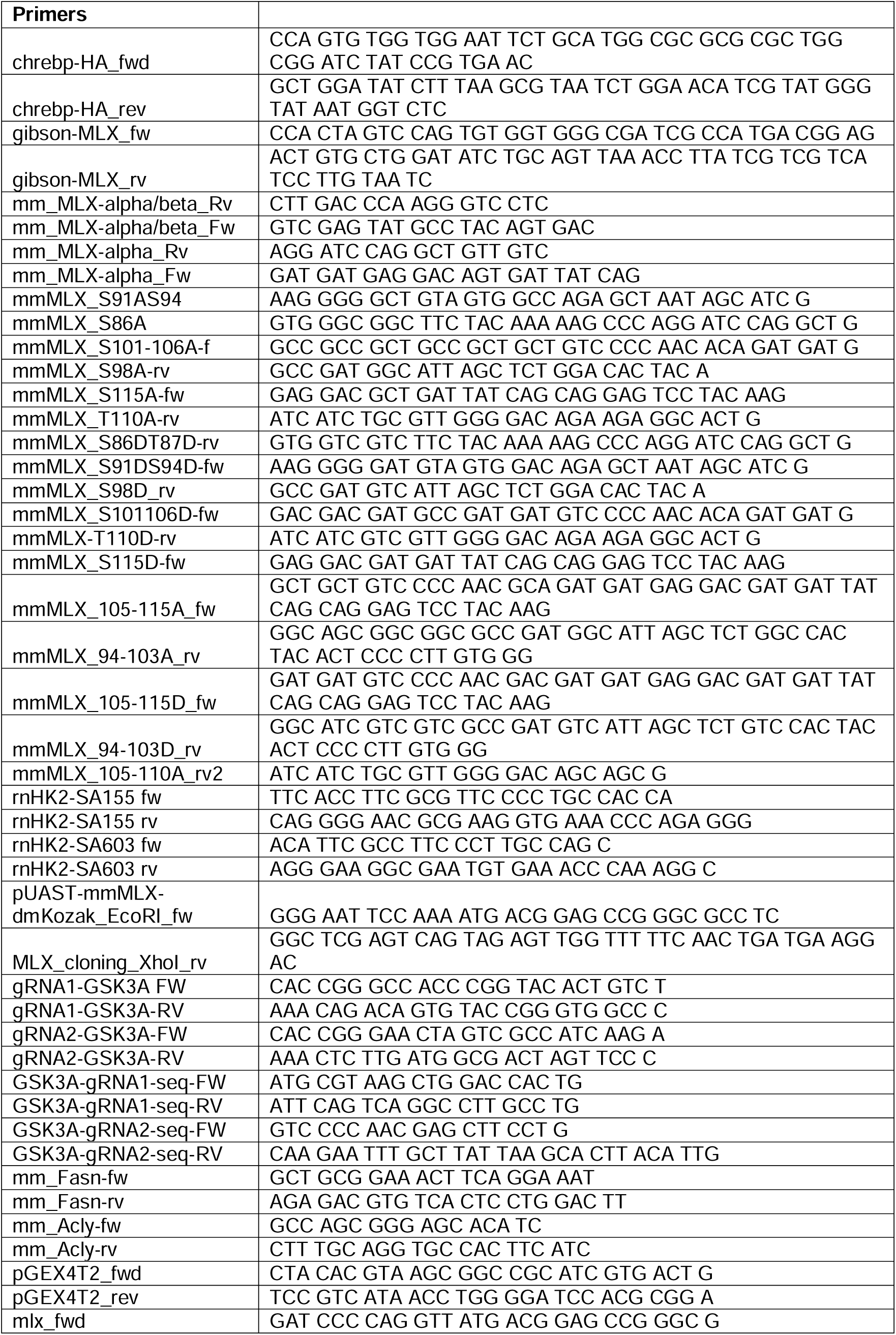
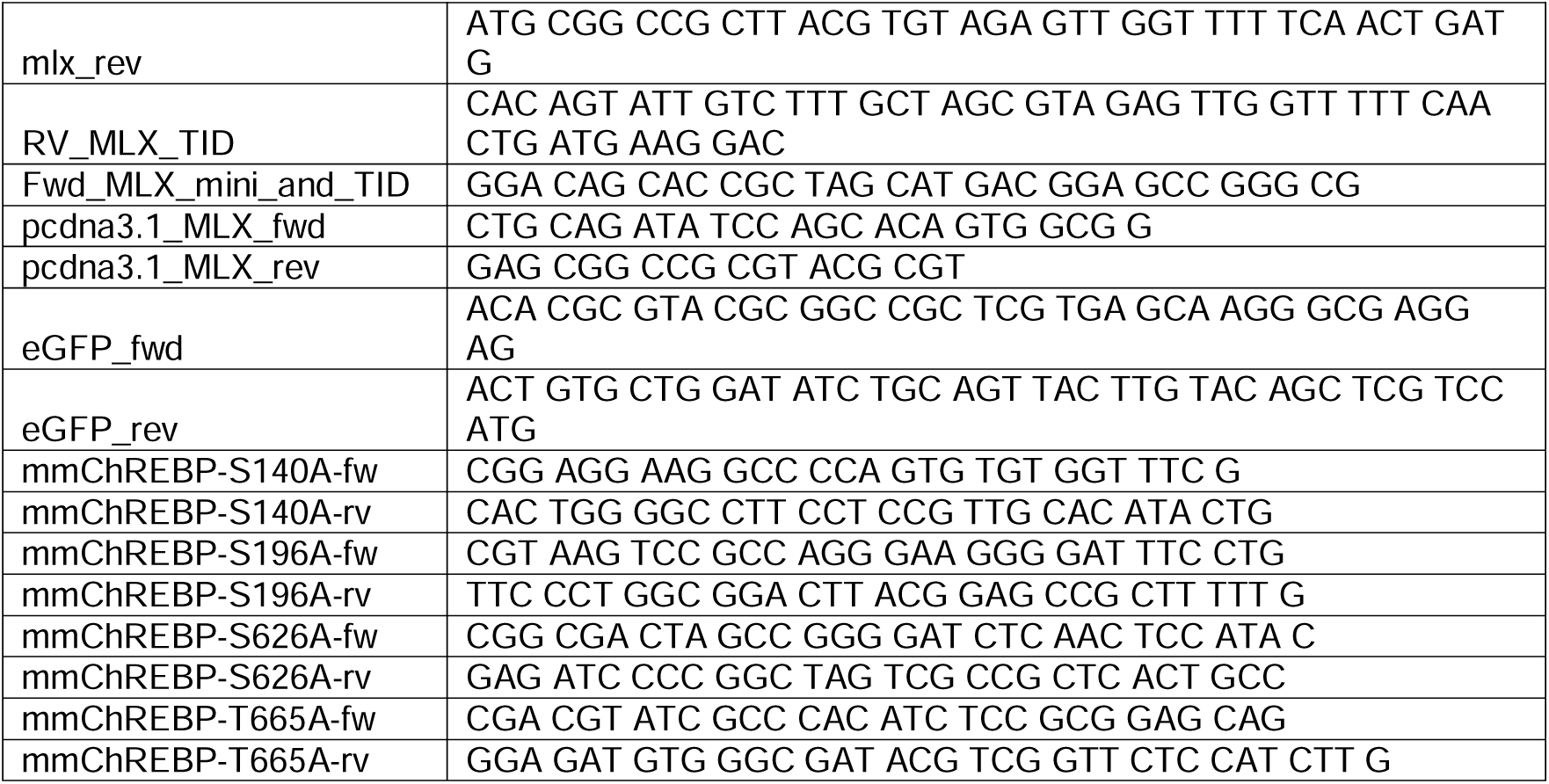

**Supplementary data table 2.**
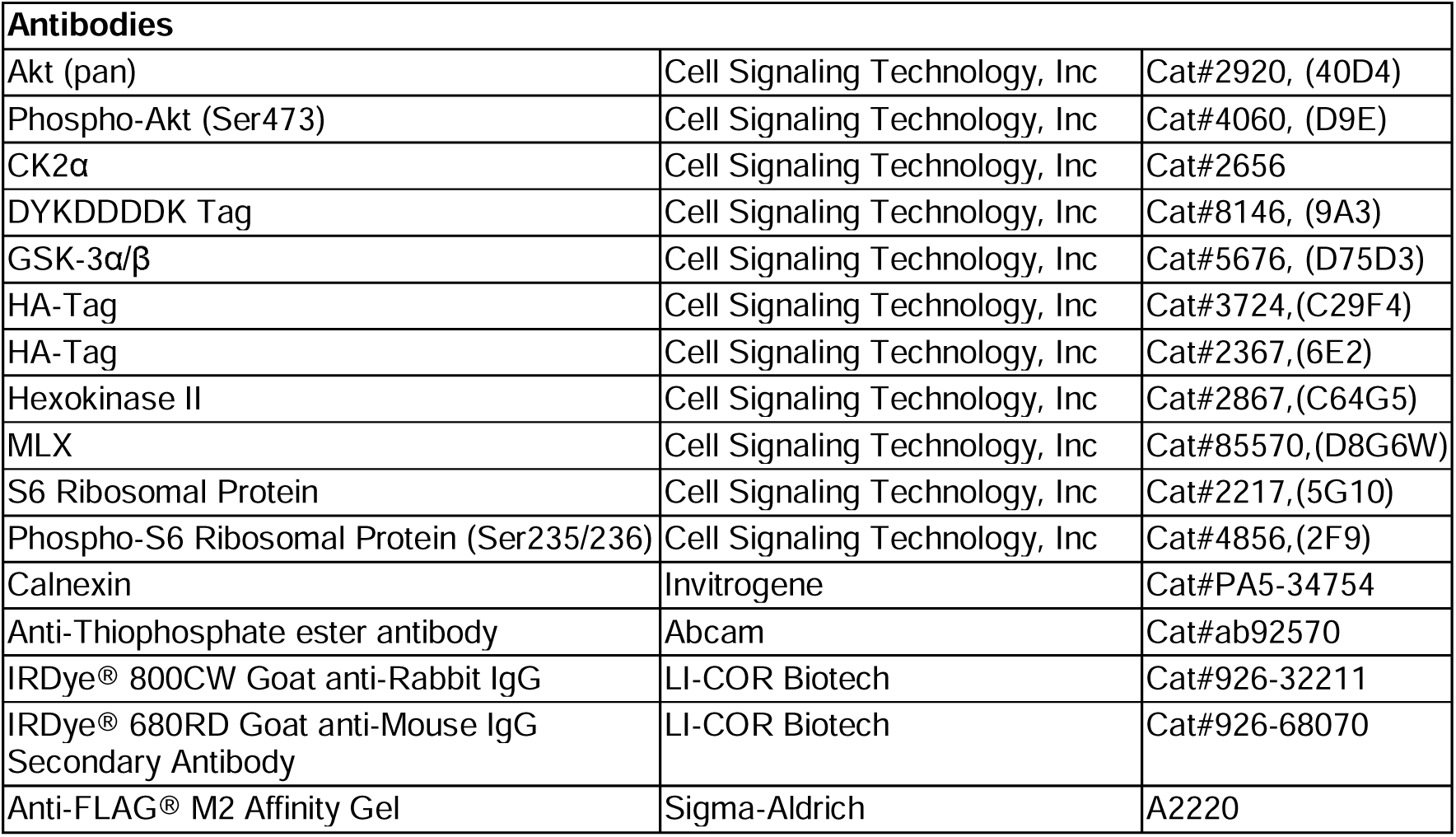

**Supplementary data table 3.**

| **Plasmids** |  |  |
| --- | --- | --- |
| pMSCV-ChREBPalpha | Michael Schupp Lab (PMID: 26181104) | N/A |
| pCMV-MLX-myc-FLAG | Origene | Cat#MR204137 |
| pCDNA 3.1 -ChREBP-HA | This paper | N/A |
| pCDNA 3.1 -MLX | This paper | N/A |
| pCDNA 3.1 -MLX-A (94-115) | This paper | N/A |
| pCDNA 3.1 -MLX-D (94-115) | This paper | N/A |
| pCDNA 3.1 -MLX-S110-115A | This paper | N/A |
| pGL3-ChoRE-luc | Michael Schupp Lab (PMID: 33023907) | N/A |
| pNL1.1[Nluc/TK] | Promega | Cat#N150A |
| pCDNA 3.1 -rnHK2 | This paper | N/A |
| Lenti-CRISPR-V2-GSK3A-1 | This paper | N/A |
| Lenti-CRISPR-V2-GSK3A-2 | This paper | N/A |
| pCDNA 3.1 -MLX-alpha | This paper | N/A |
| pCDNA 3.1 -MLX-beta | This paper | N/A |
| 3xHA-TurboID-NLS_pCDNA3 | Alice Ting Lab (Addgene) | Addgene plasmid Cat#107171 |
| pCDNA 3.1 -MLX-TurboID | This paper | N/A |
| lentiCRISPR v2 | Feng Zhang Lab (Addgene) | Addgene plasmid Cat#52961 |
| pGEX-MLX-WT | This paper | N/A |
| pGEX-MLX-WT | This paper | N/A |
| pUAST-mMLX-WT | This paper | N/A |
| pUAST-mMLX-WT | This paper | N/A |

**Figure supplementary 1.**
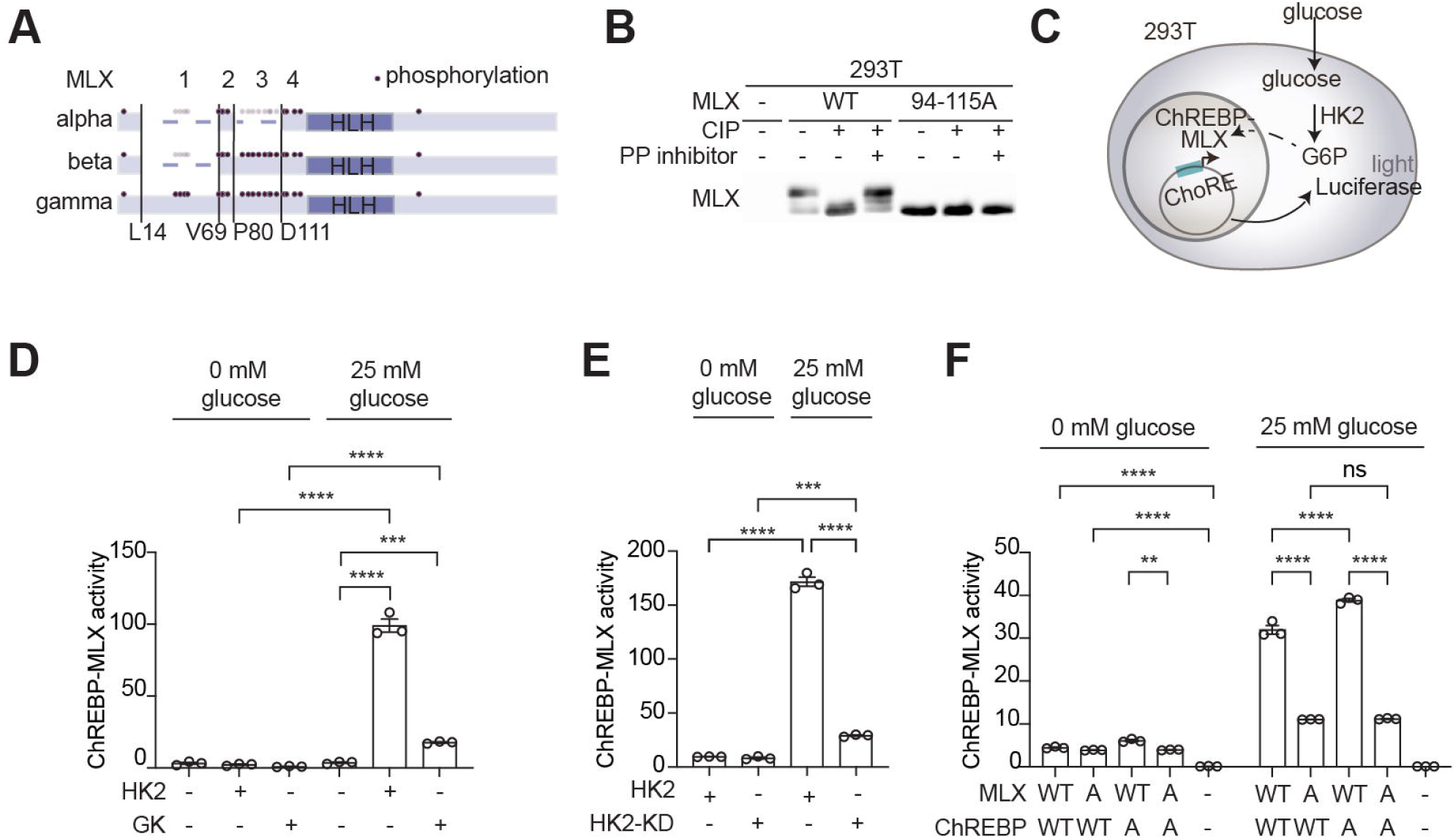
MLX phosphorylation promotes ChREBP-MLX activity, related to Fig. 1. **a.** Phosphorylation sites on MLX alpha, beta, or gamma isoforms according to PhosphoSitePlus® (https://phosphosite.org). **b.** MLX phosphorylation in 293T cells expressing MLX-WT or -A. Protein lysates were treated with calf intestine phosphatase (CIP) or CIP and phosphatase (PP) inhibitors. N=3. **c.** Luciferase reporter assay for ChREBP-MLX activity. **d.** ChREBP-MLX luciferase reporter activity in 293T expressing ChREBP, MLX-WT, and either HK2 or GCK. Cells were starved for glucose and treated with 25 mM glucose for 3 hours. One-way ANOVA, ***p<0.001, ****p<0.0001. N=3. **e.** ChREBP-MLX luciferase reporter activity in 293T cells expressing ChREBP, MLX-WT, and either HK2 or kinase dead HK2 (HK2-KD). Cells were starved for glucose and treated with 25 mM glucose for 3 hours. One-way ANOVA, ***p<0.001, ****p<0.0001. N=3. **f.** ChREBP-MLX luciferase reporter activity in 293T expressing HK2, MLX-WT or -A and either ChREBP-WT or -4A. Cells were starved for glucose and treated with 25 mM glucose for 3 hours. One-way ANOVA, **p<0.01, ****p<0.0001, ns=not significant. N=3.

**Figure supplementary 2.**
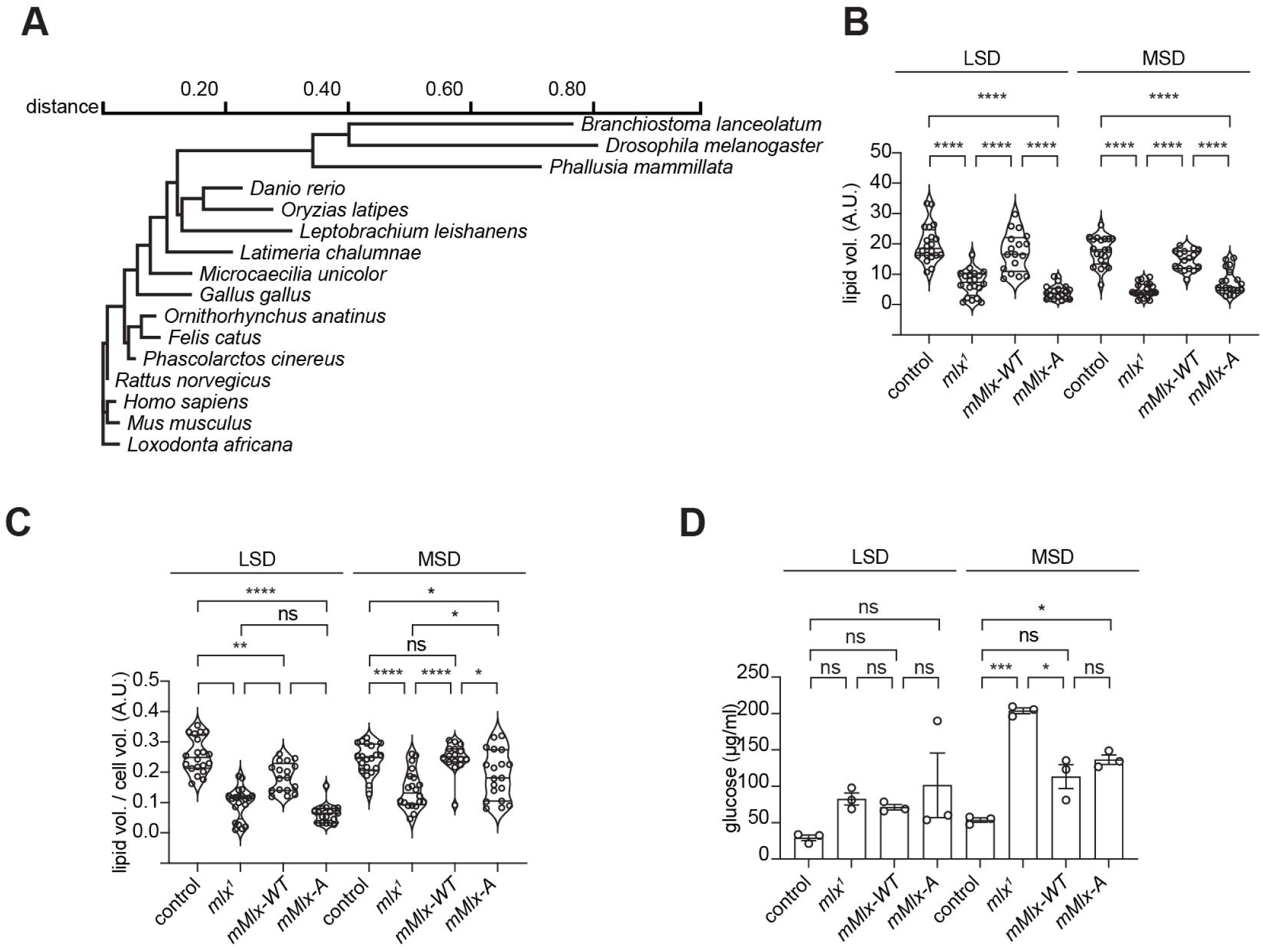
MLX phosphorylation on an evolutionarily conserved motif promotes sugar response in *Drosophila*, related to Fig. 2. **a.** Evolutionary relationships of the MLX sequence among animal species. Branching order shows the relationships between species and branch length displays the amount of evolutionary change between the nodes. **b.** Quantification of lipid staining in Fig. 2g, each point represents lipid signals per cell. Two-way ANOVA, ****p<0.0001. n=5. **c.** Quantification of lipid staining in Fig. 2g, each point represents lipid signals normalized by cell volume. Two-way ANOVA, *p<0.05, **p<0.01, ****p<0.0001, ns=not significant. n=5. **d.** Hemolymph glucose levels in 3^rd^ instar control, *mlx*^1^, *mMlx-WT* or *mMlx-A* grown in LSD or MSD. Two-way ANOVA, *p<0.05, ***p<0.001, ns=not significant. N=3.

**Figure supplementary 3.**
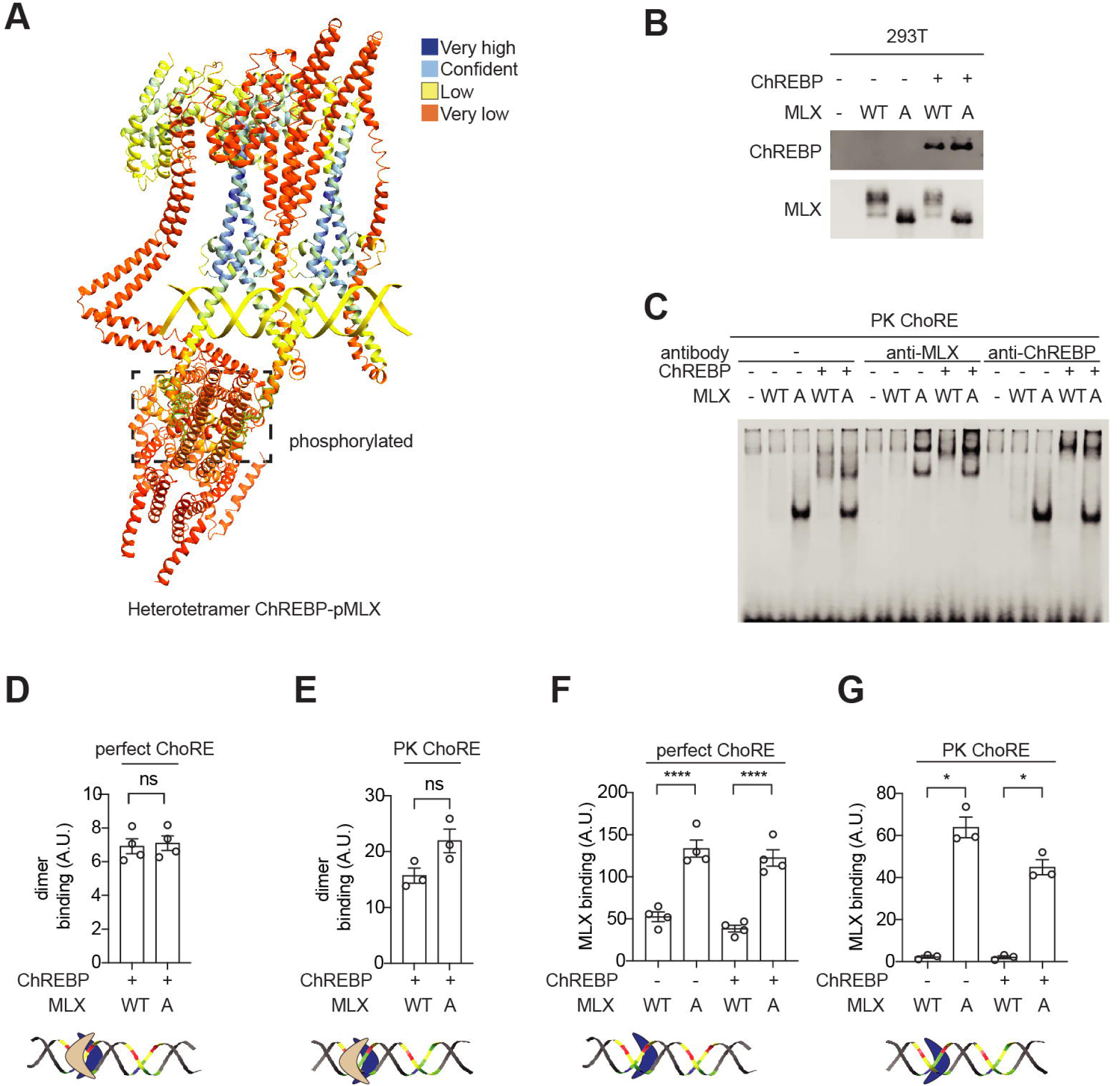
MLX phosphorylation is required for the binding of ChREBP-MLX heterotetrametric complex on the ChoRE, related to Fig. 3. **a.** Model of ChREBP-phosphorylated MLX (pMLX) structure by AlphaFold 3 with confidence scores. **b.** Immunoprecipitated MLX and ChREBP proteins from in 293T cells expressing MLX-WT or -A with or without ChREBP for Electrophoretic mobility shift assay (EMSA) experiments in Fig. 3d. **c.** EMSA with antibodies against ChREBP or MLX validating band identity from Fig. 3d and 5j. **d-g.** Quantification in Fig. 3D. t test, *p<0.05, ****p<0.0001, ns=not significant. N=3-4.

**Figure supplementary 4.**
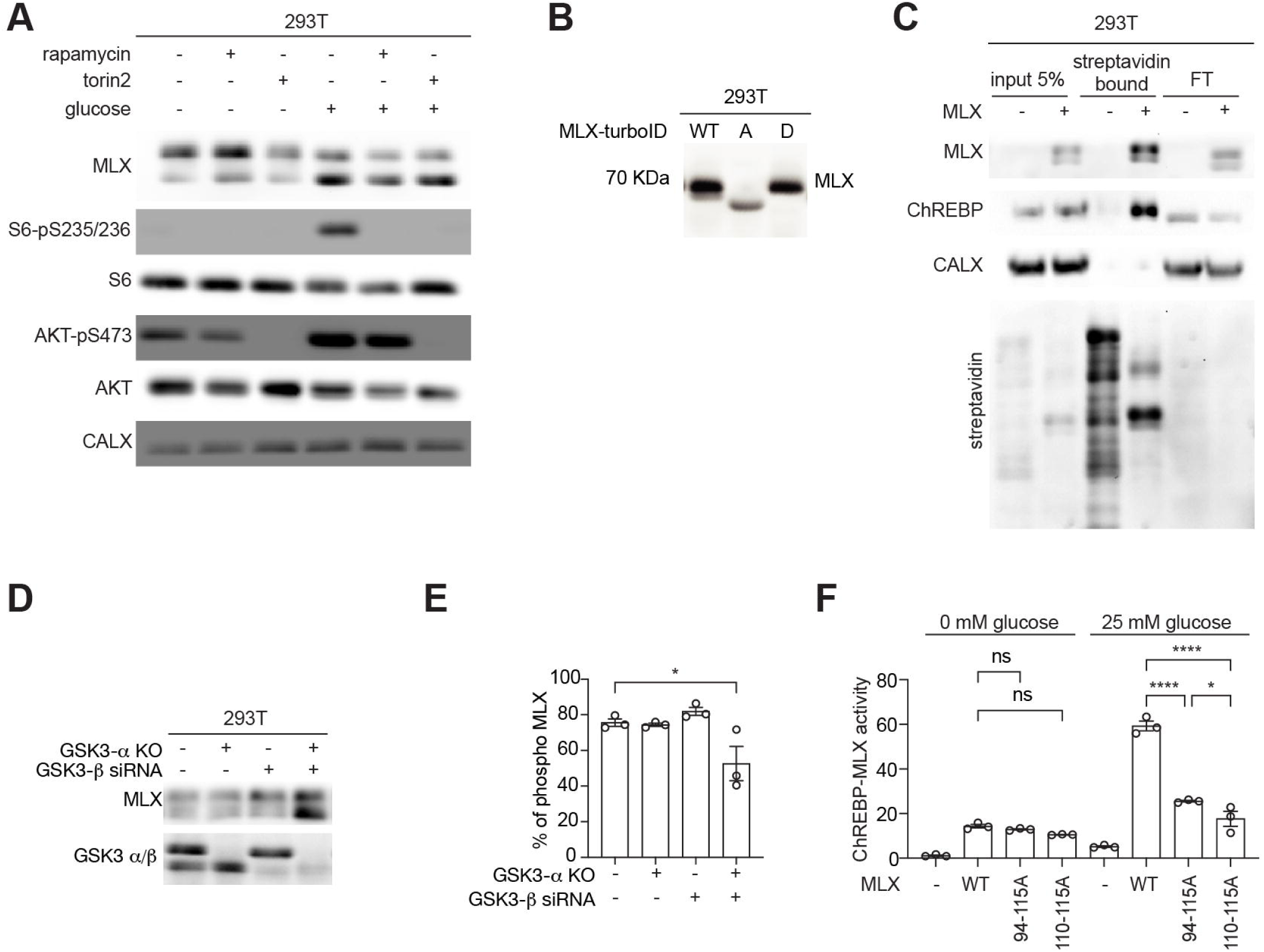
Identification of CK2 and GSK3 as MLX kinases, related to Fig. 4. **a.** MLX phosphorylation in 293T cells treated with the mTORC1 inhibitor rapamycin or mTORC1/mTORC2 dual inhibitor torin2. Cells were starved for glucose overnight, pretreated with 100 nM rapamycin or 250 nM torin2 for 30 min, and treated with 25 mM glucose for 60 min. S6-pS235/236 and AKT-pS473 serve as positive controls for mTORC1 and mTORC2 inhibition, respectively. CALX serves as a loading control. N=3. **b.** MLX phosphorylation in 293T cells expressing TurboID-tagged MLX-WT, -A and -D. N=3. **c.** Biotinylated proteins in 293T cells expressing TurboID-tagged MLX-WT, HK2 and ChREBP. Streptavidin blot serves as a control for biotinylation. N=3. **d-e**. MLX phosphorylation in WT, GSK3-a KO, GSK3-b KD or GSK3-a/b KO/KD 293T cells. GSK3-a/b serves as a control for KO/KD. One-way ANOVA, *p<0.05. N=3. **f**. ChREBP-MLX luciferase reporter activity in 293T cells expressing ChREBP and HK2 with MLX-WT, 94-115A, or 110-115A. Cells were starved for glucose overnight and treated with 25 mM glucose for 3 hours. One-way ANOVA, *p<0.05, ****p<0.0001, ns=not significant. N=3.

**Figure supplementary 5.**
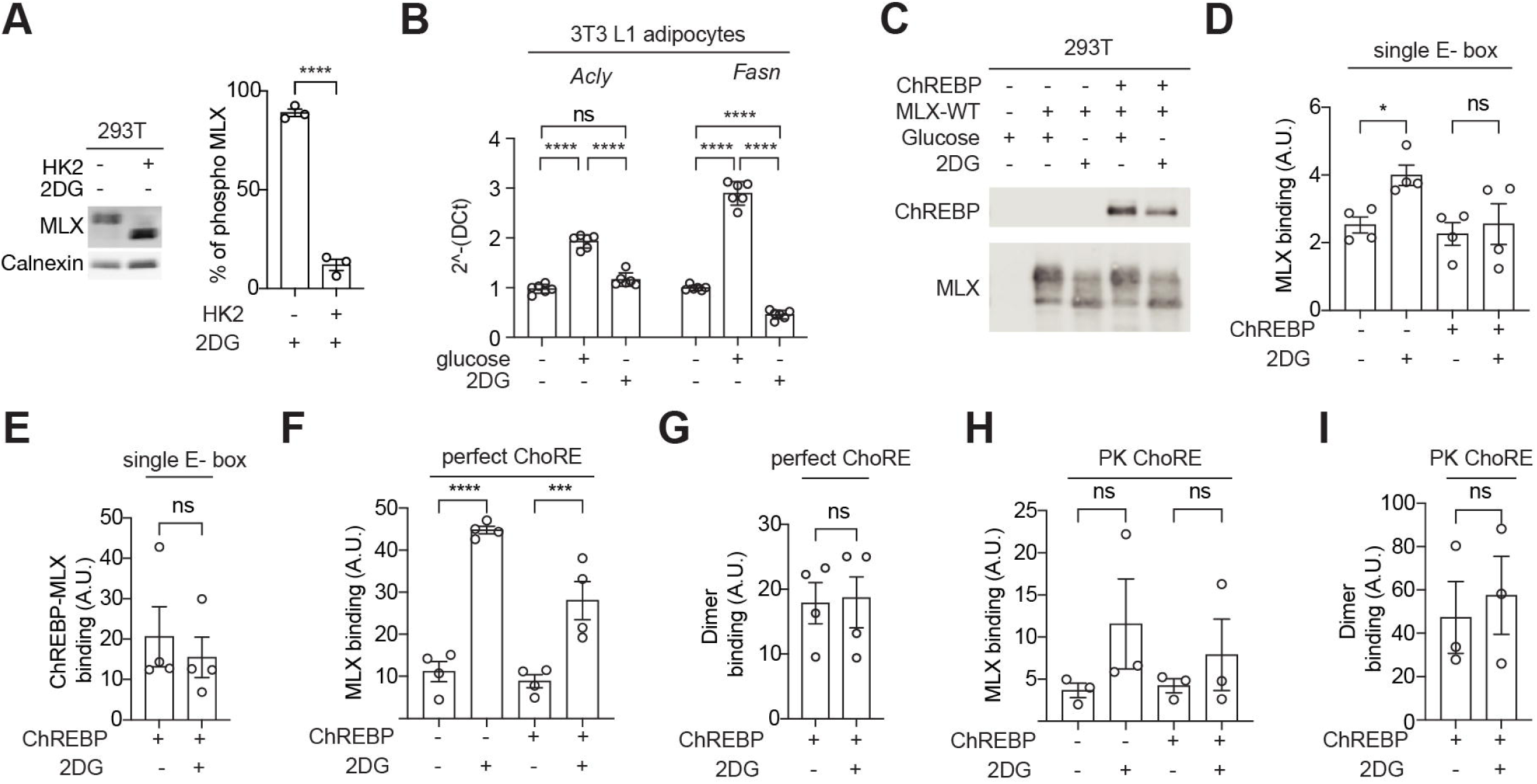
G6P accumulation inhibits CK2-mediated MLX phosphorylation and the binding of ChREBP-MLX tetramer on the ChoRE, related to Fig. 5. **a.** MLX phosphorylation in 293T cells transfected with HK2 or empty plasmid. Cells were starved for glucose overnight and refeed with 2DG for 60 min. CALX serves as a loading control. N=3. **b.** *Acly* and *Fasn* mRNA levels in 3T3 L1 adipocytes treated with 25 µM glucose or 2DG for 30 min. one-way ANOVA, ****p<0.0001, ns=not significant. N=6. **c.** Immunoprecipitated MLX and ChREBP proteins used for EMSA experiments in Fig. 5J. 293T cells expressing MLX-WT with or without ChREBP were treated with glucose or 2DG for 60 min. **d-i**. Quantification in Fig. 5J. t-test, *p<0.05, ***p<0.001, ****p<0.0001, ns=not significant. N=3-4.

## Notes

### Competing Interest Statement

The authors have declared no competing interest.

### Summary of Updates

The author order has been updated due to a mistake during the submission process

